# Factors shaping the assembly of lichen holobionts in a tropical lichen community

**DOI:** 10.1101/2024.05.29.596524

**Authors:** Magdalena Kosecka, Amélia Bourceret, Benoît Perez-Lamarque, Beata Guzow-Krzemińska, Martin Kukwa, Adam Flakus, Pamela Rodriguez-Flakus, Marc-André Selosse

## Abstract

Lichen thalli host complex microbial communities, which may foster the ecological stability and longevity of the lichen symbiosis. Yet, we lack a holistic understanding of the processes contributing to the assembly of the lichen holobiont. This study assessed the diversity and community structure in taxonomically diverse co-occurring lichens associated with Trebouxiophyceae algae from Bolivian forests. We focused on three components of the lichen holobiont: the lichenized fungus (mycobiont) and its associated algae (photobiome) and fungi (mycobiome). We specifically tested the influence of mycobiont identity, thallus morphological type, reproductive strategy, and lichen secondary metabolites on the lichen-associated photobiome and mycobiome. To understand the specialization patterns between holobiont components, we investigated interaction networks.

We observed that co-occurring mycobiont taxa host diverse, taxon-specific, yet overlapping photobiome and mycobiome. In particular, these communities are significantly influenced by the host’s thallus morphological type and its secondary metabolites. Finally, we demonstrated that both photobiome and mycobiome are structured mainly by mycobiont identity, which results in modular networks with strong phylogenetic signals and high levels of specialization. In conclusion, the symbiotic interactions within lichen are structured mainly by the mycobiont, which appears to be the leading architect of the lichen holobiont.

## Introduction

Years of studies on lichen symbiosis have repeatedly reconsidered the definition of lichens. Recently, lichens were redefined as self-sustaining ecosystems (Hawksworth & Grube 2020, 2024; Sanders 2024) formed by the main lichenized fungus (referred to as the mycobiont, which shapes the lichen thallus) interacting with one or more photosynthetic partners (referred to as the photobionts, green alga and/or cyanobacteria) as well as a microbial diversity including other algae, which together with the photobionts form the photobiome, other fungi (forming the mycobiome), bacteria (bacteriome), and viruses (virome) so that the lichen can be seen as a holobiont. These communities are more and more precisely characterized thanks to advanced DNA sequencing techniques (e.g., Spribille et al. 2016; Spribille 2018; U’Ren et al. 2019; Ponsero et al. 2021; Xu et al. 2022), and the description of the lichen holobiont have raised numerous questions regarding the interactions between its compartments within a single lichen thallus as well as interactions between these compartments in lichen communities (see Hawksworth & Grube 2020, 2024; Sanders 2024). Understanding the processes shaping the diversity and structure of the microbiome in lichen holobiont may identify factors that contribute to a stable lichen symbiosis.

The lichen mycobiome encompasses fungi that belong to three functional groups (Fernández-Mendoza et al. 2017): lichenicolous fungi (lichen parasites), endolichenic fungi (understood as inside residents of thalli), and extraneous fungi (dispersion structures of other fungi, spores, and vegetative propagules landed on the thallus) (Fig. 1a). Most lichenicolous fungi occur symptomatically (e.g., form galls) on lichen thalli, displaying their reproductive structures for spore dispersal. They are recognized as parasitic (Petrini et al. 1990; Girlanda et al. 1997; Harutyunyan et al. 2008), although they can sometimes colonize thalli without visible symptoms (Fig. 1a; Fernández-Mendoza et al. 2017). Conversely, endolichenic fungi occur inside lichen thalli as asymptomatic partners; however, their function in lichen symbiosis, from neutral to beneficial, has not yet been well defined (Fig. 1a; Arnold et al. 2009; Spribille et al. 2016; Spribille 2018). Finally, extraneous fungi, the least studied group of the lichen mycobiome, comprise accidental fungal inhabitants, including saprotrophic fungi and other species in the form of spores or vegetative propagules landed on the thallus (Fig. 1a; U’Ren et al. 2010; Muggia et al. 2010; Fernández-Mendoza et al. 2017).

**Figure 1.**
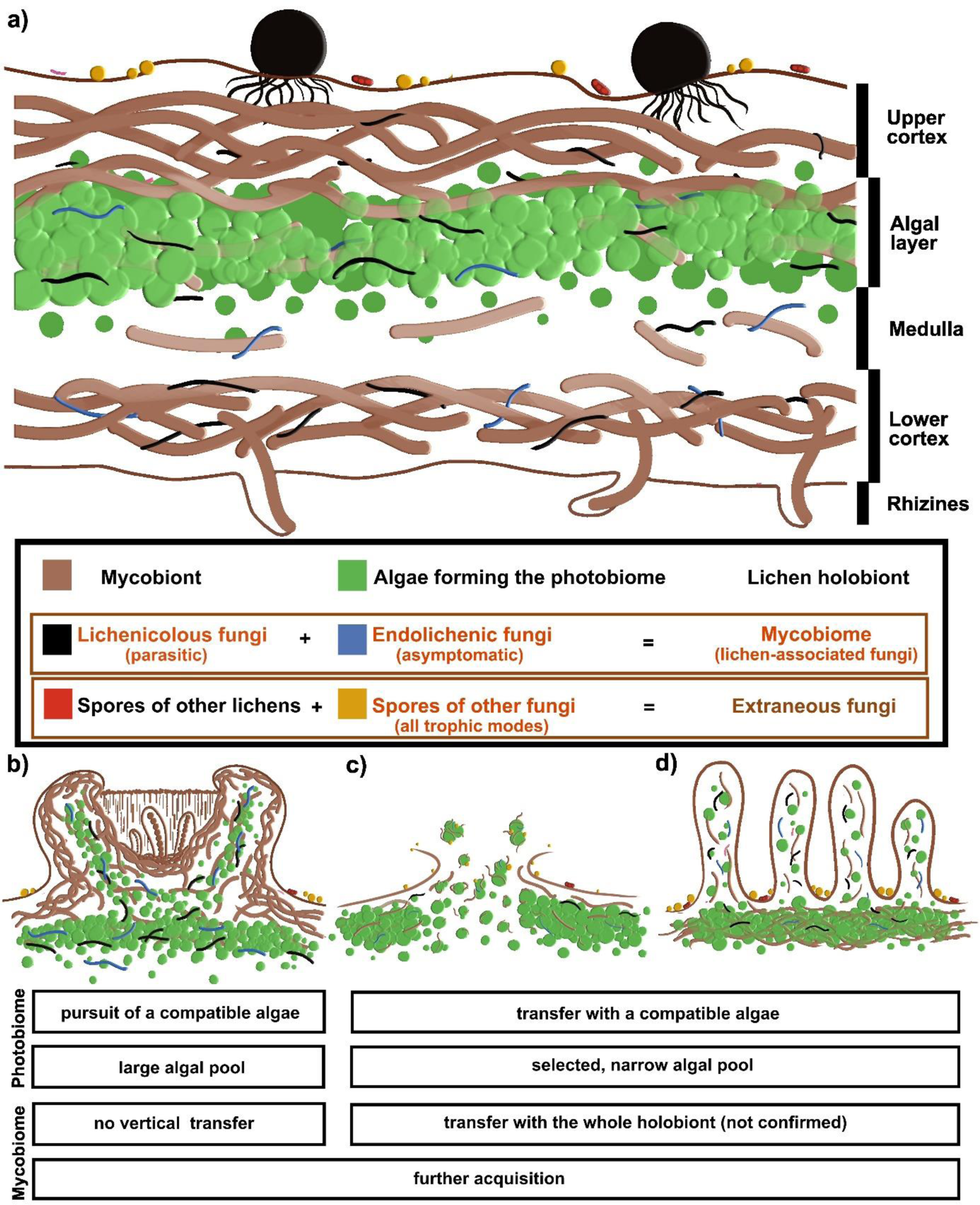
**(a)** Schematic theoretical cross-section of a lichen thallus with a heteromeric, layered organization. The scheme includes localization of the lichen-associated fungal community, also called mycobiome. It is composed of lichenicolous fungi (corresponding to pathotrophic mode), endolichenic fungi (corresponding to symbiotrophic mode), extraneous fungi that comprise the accidental fungal inhabitants that may include all trophic modes (pathotrophic, symbiotrophic, and saprotrophic) as well as other lichen spores and vegetative propagules (not shown). The scheme does not present the bacteriome and the virome, which are not the subjects of the current study. **(b-d)** Schematic theoretical cross-section of selected lichen reproduction structures: fungal apothecium (b), vegetative propagules, soredia (c), and isidia (d). The graph below the scheme shows the theoretical impact of a particular propagation mode on lichen photobiome and mycobiome.

At least three functional traits of lichens are suspected to impact their photobiome and/or mycobiome structure, namely thallus morphological type, reproductive strategies, and lichen secondary metabolites. Firstly, the thallus morphological type, which is classified into three main categories (crustose, foliose, or fruticose; Nash 2008), affects photobiome assembly. In particular, crustose lichens, which consist of a cortex layer, an algal layer, and a lower fungal layer directly and highly attached to the substrate, may be associated with a broader range of algae than foliose and fruticose taxa, whose structure is more complex, fully surrounded by a cortex layer and separated from the substrate (Fig. 1a; Helms et al. 2001). In the case of mycobiome, Stone et al. (2012) hypothesized that a high level of diversity may result from the highly porous and heterogeneous nature of lichen thalli. However, to our best knowledge, the effect of thallus morphological types on mycobiome diversity has not been addressed.

Secondly, lichen species display various reproductive strategies (Tripp & Lendemer 2018; Fig. 1b-d), which can directly impact holobiont assembly. Asexual reproduction links the mycobiont and photobiont in common vegetative propagules (dust-like soredia, Fig. 1c, or differentiated isidia, Fig. 1d), guaranteeing immediate restoration of the thallus with a compatible alga (Nash 2008), resulting in a narrower range of photosynthetic partners (Helms et al. 2001). In contrast, sexually reproducing lichenized fungi produce spores (e.g., in apothecia, Fig. 1b) that germinate before associating with compatible alga(e) to re-establish symbiosis (Nash 2008), resulting in the potential acceptance of a large range of algae (Wornik & Grube 2010; Otálora et al. 2010; Cao et al. 2015; Steinová et al. 2019). Soredia and isidia allow vertical transmission of photobiome and probably mycobiome members (Wornik & Grube 2010; Cao et al. 2015; Steinová et al. 2019; Černajová & Škaloud 2020), so that a higher specificity and a lower diversity may be expected for photobiome, and perhaps even for mycobiome. Yet, horizontal transmission during development may obscure this trend, especially for the mycobiome. The mycobiome of asexually reproducing lichens is expected to be more homogenous and specific, in contrast to sexually reproducing lichens, in which it is acquired through horizontal transmission (Arnold et al. 2009; U’Ren et al. 2010). Comparing isidia and soredia, the size of the former allows transmission of a larger fraction of the parental mycobiont community, but their cortex may limit the settlement of new fungi (especially extraneous) more than for soredia. Althogether, it is hard to predict the impact of reproduction on the mycobiome.

Thirdly, the secondary metabolites produced by the mycobiont may also modulate the communities. Lichen secondary metabolites belong to various chemical groups (Elix 2014) known for their photoprotective properties (Legouin et al. 2017; Phinney et al. 2019; Beckett et al. 2021), including resistance to high temperatures and drought (Asplund et al. 2017; Lutsak et al. 2017), allelopathic properties (Giordano et al. 1997) and antimicrobial properties (at least *in vitro*, Cohen et al. 1996; Rankovic et al. 2008). They sometimes display phytotoxicity for particular lichen-associated algae (Băckor et al. 1998, 2010; Lokajová et al. 2014), and they thus have the potential to regulate the photobiome and mycobiome.

Network-based tools constitute a powerful approach to studying interaction patterns in multispecies symbiotic systems (Dupont et al. 2003; Olesen et al. 2007). They may be useful in apprehending the assembly processes in lichen symbiosis. Most studies have revealed modular and not nested interaction networks between the mycobiont and its photobiome (e.g., Chagnon et al. 2018; Kaasalainen et al. 2021; Pérez-Ortega et al. 2023), as well as between the mycobiont and lichen-associated fungal communities (mycobiome; e.g., Chagnon et al. 2016; Yang et al. 2023). Such a network indicates the tendency of species subsets to interact preferentially and not with other species from the network. However, capturing the relationships between components of the lichen holobiont may be complicated due to the change of photosymbiont(s) depending on habitat conditions (Rodriguez et al. 2008) or the presence of many different algae within the same thalli (Paul et al. 2018). The difficulties in properly apprehending the interaction between the mycobiont and its mycobiome are due to the modulation of mycobiome by site-specific factors, the importance of which varies among mycobiont taxa (U’Ren et al. 2019). Furthermore, previous studies considered network properties of a limited assembly of lichens, i.e., one species or genera. At the scale of the complete lichen assemblage (meaning the complete multispecies lichen community), network-based tools offer the opportunity to assess how members of the holobionts are shared between lichens from different taxa.

Finally, photobiome diversity may partly structure the mycobiome (Muggia & Grube 2018). First, co-culture experiments of lichen algae and endolichenic fungi showed that the latter interact with algae by close wall-to-wall contacts and mucilage production around the contact zone and even, but rarely, form haustorium-like structures (Gorbushina et al. 2005; Arnold et al. 2009; Muggia et al. 2018). Second, endolichenic fungi are frequently present within the algal layer (Fig. 1a; Arnold et al. 2009). Thus, the interaction between the photobiome and endolichenic fungi may govern the assembly of the mycobiome.

The application of algal and fungal barcoding provides insight into the diversity of lichen holobionts. Comparison of this diversity in different co-occurring lichens with diverse thallus morphological types, reproduction strategies, and secondary metabolites offers insights into factors shaping the assembly of these complex holobionts. We hypothesize that in addition to these lichen functional traits, site conditions and host mycobiont specialization may also shape the holobiont. To assess this, we sampled co-occurring lichens with *Trebouxia* photobionts at five neotropical forest sites and investigated their mycobiome and photobiome diversities. Firstly, we tested the influence of mycobiont identity and selected lichen traits on the diversity and community structure of the mycobiome and photobiome. Secondly, we analyzed the associations between members of the lichen holobiont by reconstructing interaction networks, assessing their structures, and measuring levels of specialization between each component of the lichen holobiont. Lastly, we assessed the influence of the selected lichen functional traits listed above and site conditions on specialization levels.

## Experimental Procedures

### Sampling sites

The study was performed in three neotropical forests in Bolivia with diverse lichen assemblages, including lichens from diverse fungal groups with different thallus morphology. The dominant lichen type in selected forests are lichens that, in principle, adopt *Trebouxia* (Trebouxiophyceae, green alga) as a photosynthetic partner, while other groups that associate with Trentepohliaceae (i.e., Arthoniaceae or Graphidaceae) or with both green algae (Trebouxiophyceae) and cyanobacteria (*Nostoc*; i.e., *Lobaria*, *Lobariella*, *Sticta*) constituted a minority. Therefore, the sampling was focused only on the dominant photobiont group. Samples were collected in June 2016 in the Parque Nacional Carrasco, Department of Cochabamba (CB1-3 samplings) and Department of La Paz (LP 1-2 samplings) in the northern part of the Yungas Forest, from sites with various levels of anthropogenic disturbance. The sampling site CB1 was partly anthropogenically disturbed (especially grazed by cattle and with tree logging), whereas the site CB2 was minimally influenced by human activities (Table S1). These two sites were located at similar altitudes and separated by a distance of ca. 400 m. From each site, 15 *Alnus acuminata* trees (the dominant species) were selected, and 360 lichen specimens were collected in total. CB3, placed ca. 8 km away from CB1 and 2, was minimally influenced by human activities and by tree logging while retaining similar lichen assemblages: 28 specimens were harvested from *Alnus acuminata*. The two other sites were located in the neighboring Department of La Paz in a savanna-like ecosystem (Table S1): LP1 was undisturbed while LP2 was more disturbed (grazed by cattle and with tree logging), and 53 and 35 specimens were respectively sampled from the same host tree as above. All samples were frozen until further procedures.

The morphology and anatomy of lichen thalli were used to assess the taxonomy to the genus level. Thin-layer chromatography (TLC) was used to identify the lichen secondary metabolites necessary for the recognition of many groups of lichens, according to Culberson and Kristinsson (1970) and Orange et al. (2001). To study the influence of secondary metabolites on the lichen holobionts, we grouped the identified metabolites into 12 substance classes and coded their presence in the specimen by 1 and their lack by 0 (Table S2).

### Microbial community profiling

A small piece of thallus was used without surface sterilization from each lichen specimen to get a complete lichen holobiont. Libraries were prepared according to Bourceret et al. (2022). Samples were processed in triplicate by polymerase chain reaction (PCR) using specific primer sets for the profiling of fungal communities, comprising the main mycobionts and lichen-associated mycobiome, and algal communities that form the photobiome, comprising the photobionts as well as surface algal inhabitants (internal transcribed spacer 2 [ITS2], see PCR conditions in Methods S1). Multiplexing was done using a unique pair of barcoded primers in the PCR, following Petrolli et al. (2021). Two libraries were prepared in equimolar amounts to build one library per group (fungi and algae; Methods S1). Sequencing was performed using a MiSeq, paired-end 2*250bp, Illumina technology (Fasteris, Switzerland).

### Amplicon sequencing data processing

Sequencing outputs from the two libraries (the ‘fungal dataset’ and ‘algal dataset’) were separately processed using VSEARCH (Rognes et al. 2016) following Perez-Lamarque et al. (2022a). In short, paired-end reads were assembled, quality checked, and demultiplexed with cutadapt (Martin 2011). Fungal reads were clustered into 97% operational taxonomic units (OTUs), while algal reads were processed into amplicon sequence variants (ASVs) following the recommendations of Blázquez et al. (2022). Clustering algal reads into 97% OTUs provided qualitatively similar results, therefore only results on ASVs are reported here. OTUs or ASVs were checked for chimeras. For fungal taxonomic assignments, we used UNITE (Nilsson et al. 2019) and our database composed of reference sequences of fungal mycobionts obtained using Sanger sequencing (unpublished data). For algae, we used Silva (Quast et al. 2013) amended with sequences obtained by Kosecka et al. (2022). Contaminants were filtered out of the OTU/ASV tables using the decontam pipeline (Davis et al. 2018). We removed OTU/ASV present in less than 5 reads and converted the abundances into relative abundance while only retaining OTU/ASV representing at least 1% of the reads in each sample (Perez-Lamarque et al. 2022b).

We prepared three different OTU/ASV tables that described different compartments of the lichen holobiont: the mycobiont, the photobiome, and the mycobiome (including all lichen-associated fungi: lichenicolous, endolichenic, and extraneous fungi, but excluding the mycobiont(s); see Methods S2). To identify the category for each mycobiome’s OTU, we first used FUNGuild (Nguyen et al. 2016) to assign a guild, i.e., pathotroph, symbiotroph, or saprotroph. This was considered as a first basis for assignment to, respectively, lichenicolous, endolichenic, and extraneous fungi, validated after expert visual checking (e.g., removing animal or plant pathotrophs from extraneous fungi). This approach may limit the exactness of the delineation of extraneous fungi, which may contain OTUs that we attribute to other guilds. However, the exact delimitation of the location and role of a particular fungus is beyond the scope of the present research. All of the following analyses were performed on R (R Core Team 2023).

### Diversity analyses

We first investigated differences in the community composition of the photobiome and the mycobiome according to four factors: (i) the mycobiont identity at the order and genus level; (ii) the thallus morphological type; (iii) the lichen reproductive strategies; and (iv) the sampling site (Table S2). We computed beta diversities using the Bray-Curtis index with the R-package *vegan* (Oksanen et al. 2012) and performed principal coordinate analysis (PCoA). To test for significant differences across groups, we used PERMANOVA (*capscale* and *anova.cca* in *vegan* package). We also tested whether lichens with similar secondary metabolites tend to host similar photobiomes or mycobiomes using Mantel tests (*mantel.test* in *ape* package) (Paradis & Schliep 2019).

Second, we tested the variations of alpha diversities of photobiome and mycobiome communities within lichens according to the same four factors and the presence of secondary metabolite groups. Alpha diversities were computed using either observed OTU/ASV richness or the Shannon index with *vegan* (Oksanen et al. 2012). Differences between groups were tested using Kruskal-Wallis tests (*kruskal.test* function). We additionally performed rarefaction analyses on the observed richness (*rarefy* in *vegan* package) (Oksanen et al. 2012), which indicated that sites LP1-2 and CB3 were far from reaching saturation, while sites CB1 and 2 were closer to reaching a plateau (Fig. S1). We therefore only kept the two well-sampled communities (CB1 and 2) for all the following analyses, which may be sensitive to undersampling.

Finally, we investigated whether the assembly of the mycobiome was primarily influenced by the mycobiont and/or the photobiome composition. We partitioned the variance observed in the Bray-Curtis dissimilarities of the mycobiome against the Bray-Curtis dissimilarities of the mycobiont or the photobiome using *varpart* in the *vegan* package (Oksanen et al. 2012). We used ANOVA to test whether using both mycobiont and photobiome dissimilarities explains the variance observed in the mycobiome better than using mycobiont or photobiome dissimilarities alone. We performed this analysis within each site, focusing on all lichens or separately looking at the dominating lichen genera (*Usnea, Parmotrema, Hypotrachyna, Polyblastidium,* and *Leucodermia*). We also replicated the analyses using UniFrac dissimilarities instead.

### Reconstructing interaction networks and assessing their structures and specializations

We assessed the patterns of interaction within the lichen holobionts. Based on the OTU tables, at sites CB1 and CB2, we reconstructed three types of bipartite interaction networks: (i) the network between the mycobiont and the photobiome, (ii) the network between the mycobiont and the mycobiome, and (iii) the networks between the photobiome and the mycobiome. We built quantified OTU-level networks: for instance, for the mycobiont-photobiome network, for each pair of mycobiont OTU and algal ASV, we counted the number of samples where both taxa were present.

We evaluated the structure of each network using different metrics following Perez-Lamarque et al. (2022b). First, we computed the connectance as the ratio of observed interactions within each network (Dormann et al. 2008). Second, we computed the nestedness, i.e., the tendency of specialist taxa to interact with generalist taxa, which forms an interaction core, using the *nested* function (with the weighted NODF method) (Almeida-Neto et al. 2008). Third, we computed modularity, i.e., the tendency of subsets of taxa to preferentially interact with each other, using the function *computeModules* (Dormann et al. 2008). Fourth, we measured the network-level H2’ index of interaction specialization (*H2fun*) (Dormann et al. 2008), which indicates whether interactions within the network tend to be random (H2’ value close to 0) or highly specialized between partners (H2’ value close to 1). We also investigated the levels of specialization of each OTU/ASV using the net relatedness index (NRI) metric: for each taxon, NRI quantifies the mean pairwise phylogenetic distance between all its partners - a low NRI value indicates that the taxon tends to be highly specialized toward closely related partners (Webb et al. 2002). Finally, we measured phylogenetic signals in species interactions within each network using Mantel tests (Perez-Lamarque et al. 2023).

To assess the significance of the different indices (connectance, nestedness, modularity, and network-level index of specialization), we compared the original index values to the distribution under a null hypothesis obtained by shuffling the OTU tables before reconstructing the networks. For this purpose, we shuffled 100 times independently the sample names in the OTU tables, built 100 randomized networks, and computed the indices again.

## Results

### The photobiome and mycobiome composition

From 309 lichen specimens that succeeded in the amplification of the mycobiont, photobiome, and mycobiome (Fig. 2), we retrieved 111 main mycobiont OTUs, 183 algal ASVs, corresponding to the photobiome, and 1100 non-lichenized fungal OTUs, corresponding to the mycobiome. The majority of algae detected belong to the genera *Trebouxia* (Fig. 2b), except for Teloschistales mycobionts, whose photobiome is mainly formed by species of *Symbiochloris* (another genus within Trebouxiophyceae; Fig. 2b). In the mycobiome community profiles, we observed that in general species from the Caliciales and the Lecanorales have similar mycobiomes and that the species from the Pertusariales and the Teloschistales have mycobiomes respectively dominated by Mycosphaeralles and Asterinales (Fig. 2e).

**Figure 2.**
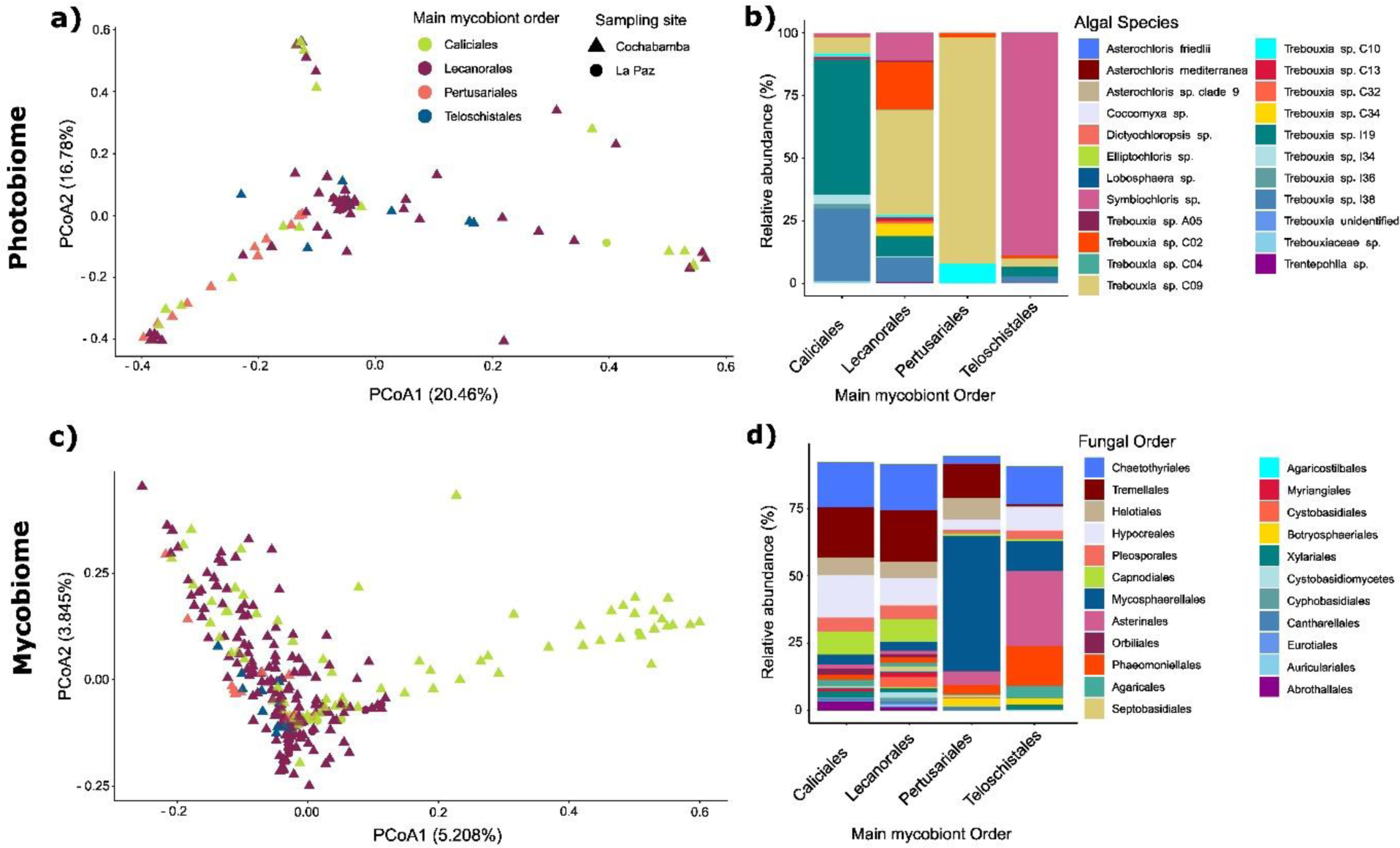
The main mycobiont identity at the genus level shapes both lichen-associated photobiome (a,b) and mycobiome (c,d). Relative abundances of the most abundant algal (b) and fungal (d) taxonomic groups are displayed at the species and order levels, respectively. PCoA based on Bray-Curtis dissimilarity (a, c) was performed for all specimens (n = 309), and shapes indicate the sampling site (Cochabamba or La Paz), whereas colors present the main mycobiont identities at the genus level. PERMANOVA revealed significant differences within mycobiont orders in both photobiome (R^2^=0.26, p=0.001) and mycobiome structure (R^2^=0.15, p=0.001).

### Factors shaping the photobiome and mycobiome composition

We tested the influence of the mycobiont identity (order and genus level), the thallus morphological type, the lichen reproductive strategies, and the sampling site on the composition of the photobiome and mycobiome. The mycobiont identity at the order or genus level appeared to be the leading source of variation of community profiles for the photobiome and mycobiome (Fig. 2 & Fig. S2B, S2D), as well as the relative abundance of all fungal trophic modes within the mycobiome, i.e., lichenicolous, endolichenic and extraneous (Fig. S3). PCoA confirmed that the mycobiont taxonomic assignment (order level) is a principal driver for both the photobiome (Fig. S2A) and mycobiome community structure (Fig. S2C). The mycobiomes were clustered on the first axis (5.2% of variance) according to the Lecanorales, Pertusariales, and Teloschistales mycobiont orders, whereas samples assigned to the Caliciales mycobiont were more dispersed (Fig. S2C). Similarly, we observed a slight clustering of samples according to the mycobiont genus (ex. *Hypotrachyna, Usnea*, and *Parmotrema*) (Fig. S2C). This trend was a bit less pronounced for the photobiome at the mycobiont order level (Fig. S2A) and genus level (Fig. 2a). PERMANOVA analyses further confirmed a significantly stronger effect of the mycobiont genus level on the photobiome and mycobiome community structure (Table S3; R2: 0.26, 0.15, respectively) than the order level (Table S3; R2: 0.11, 0.05).

By partitioning the variance between mycobiomes of different lichens, based on the observed variations of the mycobionts and photobiomes, we revealed a significant contribution of the mycobiont (Fig. S4; 13%, 8%, and 12% at all the sites, CB1 and CB2, respectively; ANOVA: p<0.05), but no contribution of photobiome alone (−3%, −5%, and −3%) to the mycobiome composition. Although we detected a potential joint contribution of the mycobiont and photobiome to the mycobiome composition (7%, 8%, and 6%), we cannot exclude that photobiome composition has no significant effect (ANOVA: p>0.05; Table S5). We observed similar results when considering the lichenicolous or endolichenic fungi (Fig. 3), but not for the extraneous fungal community, which was not significantly affected by either the mycobiont or photobiome (Fig. 3c). These results also hold when applied at the level of the mycobiont genera (Table S4).

**Figure 3.**
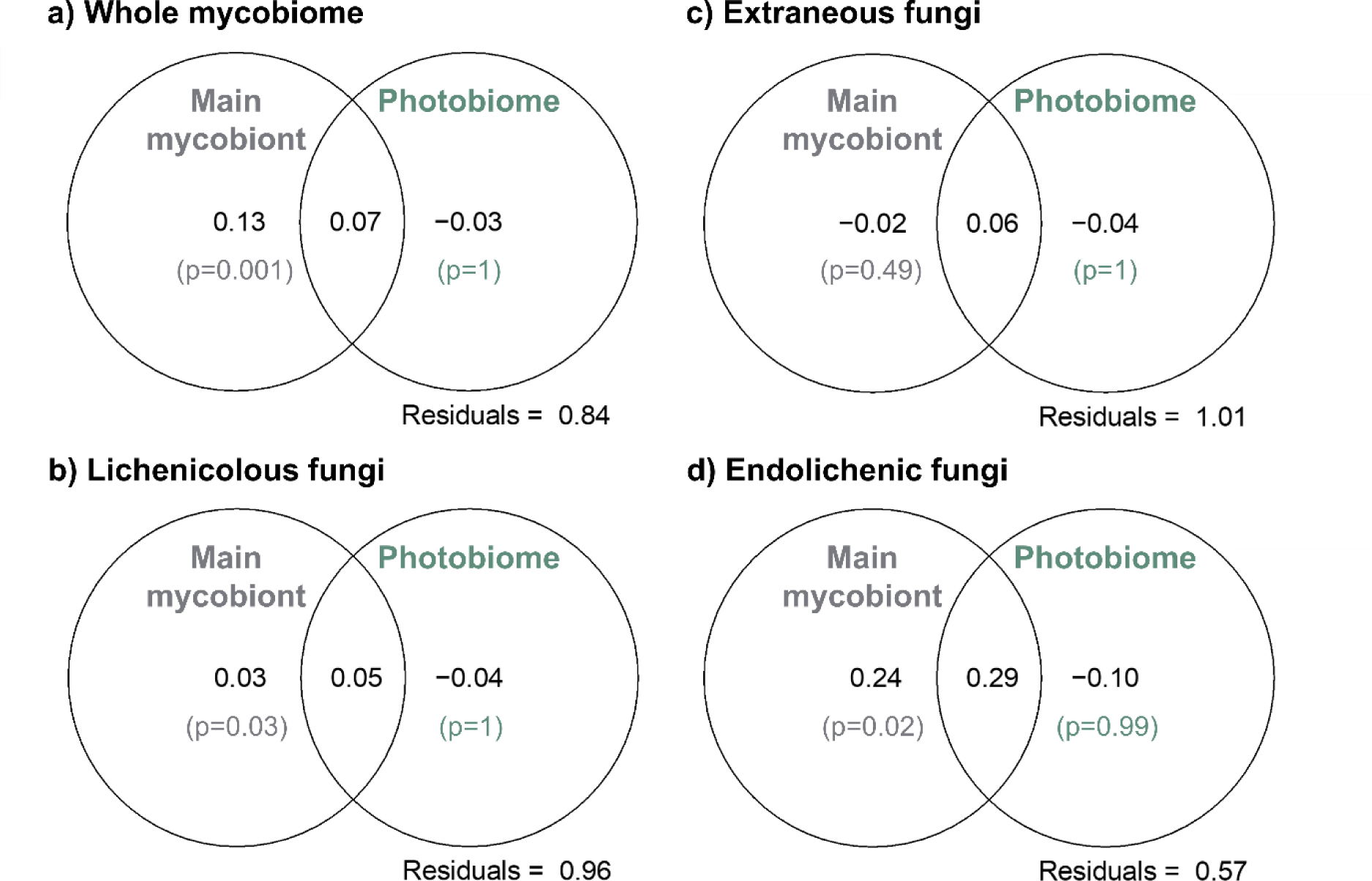
Variance partitioning indicated that the main mycobionts, but not the photobiome, have the main impact on the lichen-associated mycobiome, i.e., all fungi forming the mycobiome (a), or only lichenicolous (b), extraneous (c), or endolichenic (d) fungi according to FUNGuild. We partitioned the variance between mycobiomes as a function of the dissimilarities in the main mycobionts and photobiome communities (all measured using Bray-Curtis dissimilarities or weighted UniFrac distances). Using Venn diagrams, we reported the percentage variance that is explained by the main mycobionts or the photobiome communities or by a combined effect of both communities. The p-values (p) indicate the effects of the main mycobionts or the photobiome assessed using ANOVA (with 999 permutations). Significance results were identical when using weighted UniFrac distances.

Thallus morphological type significantly affected photobiome and mycobiome composition (Fig. 4). Yet, PERMANOVA analyses indicated a lesser effect than that of the mycobiont identity (Table S3). Thallus morphological type also significantly influenced the richness of the photobiome (number of ASVs) and mycobiome (number of OTUs) (Fig. 4a, 4b, Table S3). For the photobiome, the number of ASVs (Fig. 4a) and the Shannon index (Fig. S5a) were significantly increased in crustose lichens, regardless of the identity of the mycobiont. For the mycobiome, the number of OTUs was significantly increased in crustose and foliose Lecanorales (Fig. 4b), compared with the fruticose representatives, but the effect was not observed beside this order.

**Figure 4.**
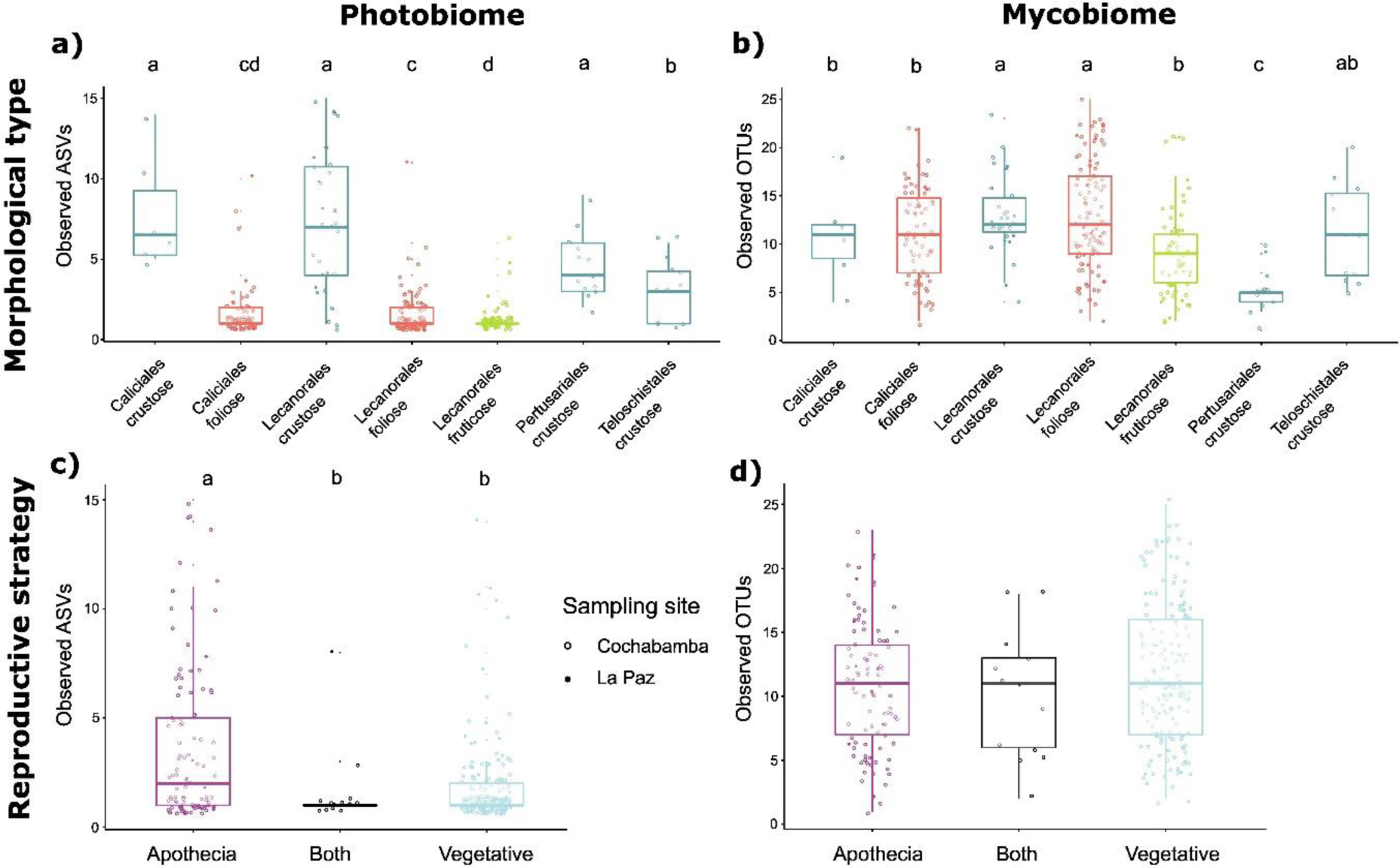
Thallus morphological type (a,b) and reproductive strategies (c,d) shape the alpha diversity of lichen-associated photobiome (a,c) and mycobiome (b,d). Letters indicate significant differences between conditions (p < 0.05) for a Kruskal-Wallis test. For reproductive strategies, “apothecia” corresponds to lichens that form only apothecia; “vegetative” corresponds to lichens with one or two vegetative propagule types (isidia or soredia), and “both” to lichens with apothecia and one or both vegetative propagule types.

The reproductive strategy significantly affected photobiome richness (Table S3). Lichens that form only apothecia (fungal reproduction sexual spores) displayed an increased number of ASVs (Fig. 4c, S5A) and higher Shannon index (Fig. S4c, S6c) compared to lichens with isidia and/or soredia (asexual reproductive strategy linking the two partners). We found no difference in photobiome between lichens with isidia vs. soredia (Fig. S6a, c). For the mycobiome, we observed no significant difference in the number of OTUs, in the Shannon index, or between lichens with apothecia vs. those with isidia and/or soredia (Fig. 4d, S5d). However, within asexually reproducing lichens, we observed higher OTUs abundance in the mycobiome of those with isidia vs. those with soredia (Fig. S6a).

Considering spatial variations, we observed small differences in community profiles and alpha diversities (Fig. S7) between the two most (and equally, n=261) sampled sites, but these differences were not significant. Rarefaction curves indicate that the undisturbed site (CB2) exhibited a higher richness of mycobionts, photobiome, and mycobiome than the disturbed site (CB1; Fig. S1). The sampling site had an overall negligible effect on community compositions (Table S3; PERMANOVA, R2<0.05): dominating mycobionts, photobiome, and mycobiome at both sites have a similar identity, which indicates the retention of the same communities.

Lichens with similar secondary metabolite composition (Table S2) tended to host similar photobiome and mycobiome communities (Mantel tests; Table S5a). In particular, beta-orcinol depsides and terpenoids had the most significant effect on the photobiome and mycobiome communities, which was observed in all lichen specimens in which these secondary metabolite groups were identified (Table S5b): in particular, these two were associated with a larger proportion of extraneous and endolichenic fungi (Wilcoxon tests: p<0.01). Conversely, orcinol depsides were associated with an increased proportion of lichenicolous fungi (W=1111, p=0.02). In terms of alpha diversity, lichens devoid of secondary metabolites displayed, on average, a more diverse photobiome (Wilcoxon test: W=382, p=0.03), while those without beta orcinol depsides were colonized by a more diverse mycobiome (W=8269, p=0.003). Lichens with orcinol depsides simultaneously hosted a richer photobiome and mycobiome (Wilcoxon tests: p<0.05).

### Network structures and specialization

The direct or indirect interactions within lichens resulted in contrasted network structures and different levels of specialization (based on the communities retrieved from two sites, CB1 and CB2; Fig. S8a, S9). The networks linking mycobiont and algae forming the photobiome appeared to be significantly modular, non-nested, and presented a significant level of specialization, indicated by a lower connectance (0.05) and higher H2’ values (H2’>0.20) than expected under the null model (Fig. 5, S8a & S9). These networks also presented significant phylogenetic signals on both sides, indicating that phylogenetically related mycobionts tend to interact with similar algae in the photobiome and *vice versa* (Table S6), as also suggested by the trend of genus-level clustering in the network representation (Fig. 5a). Considering OTU-level (or ASV-level) specialization individually, we observed a significant net relatedness index (NRI) for some mycobionts, but not for photobiome, suggesting that these mycobiont OTUs tend to associate with phylogenetically clustered algae, but not *vice versa –* i.e., a given alga may associate with different mycobionts (Fig. S10). In other words, closely related mycobionts tend to interact with closely related algae, but closely related algae tend to interact with similar, but not necessarily closely related mycobionts. Specialist mycobionts were particularly abundant among *Megalaria* and *Pertusaria* (Fig. S10a).

**Figure 5.**
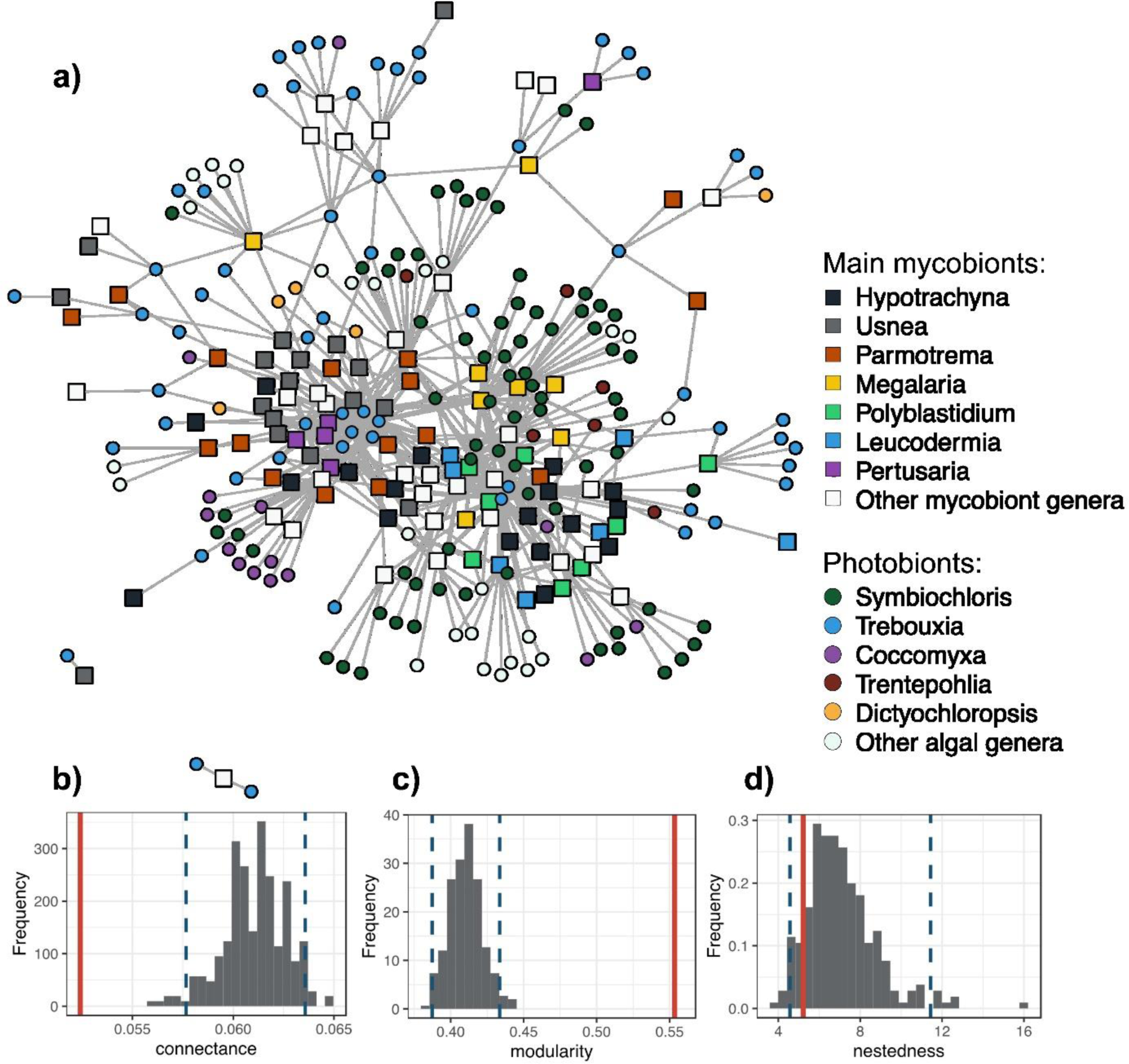
Mycobiont-algae networks exhibit a modular, non-nested, and phylogenetically conserved structure. **(a)** Representation of the meta-network between all mycobionts and algae across the 309 lichens collected across 5 sites. Each node represents a mycobiont OTU (square shape) or an algal ASV (circular shape), and each gray line indicates an interaction. The position of the nodes reflects the similarity in species interactions using the Fruchterman-Reingold layout algorithm from the igraph R-package. OTUs and ASVs are colored according to their genus. **(b-d)** Evaluation of the structure of the mycobiont-algae network reconstructed in an undisturbed community of Cochabamba (CB2): we measured the original connectance (b), modularity (c), and nestedness (d) values (represented in red) that we compared to the null expectations given by randomized networks (in gray). Dashed lines delimit the estimated 95% confidence intervals.

The networks formed between mycobionts and their mycobiome were also significantly modular, non-nested, and presented a significant level of specialization (based on both connectance and H2’; Fig. S8b, S9). However, the level of specialization (H2’ between 0.12 and 0.20) was lower than that between mycobionts and algae (H2’>0.20). Furthermore, we detected limited evidence of OTU-level specialization (Fig. S10b) or phylogenetic signal (Table S7), especially at the disturbed site (CB1). Also, we found significant differences between the ecological guilds of lichen-associated mycobiome: endolichenic fungi tended to be more specialized in their interactions than extraneous and lichenicolous fungi (Fig. S11).

Similarly, we also observed modular and non-nested networks in the case of interactions between photobiome and mycobiome (Fig. S8c). However, they presented significant but low levels of specialization (H2’<0.12; Fig. S9), intermediate levels of OTU-level specialization (Fig. S10c), and moderate evidence of phylogenetic signal (Table S7). Levels of specialization (H2’) tended to be higher in foliose lichens in comparison to crustose and fruticose morphological types, in terms of mycobiont-photobiome interactions (H2’>0.4; Fig. S12a), and lower (H2’</=0.2; Fig. S12a) in terms of mycobiont-mycobiome interaction.

Propagation modes also affected mycobiont-photobiome specialization as H2’ values were lower in lichens forming apothecia than in lichens reproducing vegetatively (Fig. S12b). Finally, network structure differed slightly between the two studied sites (CB1 and CB2): H2’ values in networks between mycobionts and photobiome as well as between mycobionts and mycobiome were lower at the disturbed site (CB1) than at the undisturbed site (CB2; Fig. S9, S12a, S12b & S13; LP1 and LP2 were not compared because the dataset was too limited). Furthermore, considering the thallus morphological type in this estimation, the above pattern was observed in foliose and fruticose, but not in crustose lichens (Fig. S12a).

## Discussion

Using the combination of functional traits and network-based analyses, we revealed that the assembly of lichen holobionts is influenced by mycobiont identity, thallus structure, reproductive strategies, and lichen secondary metabolites. Importantly, mycobionts, not algae forming the photobiome, impact the mycobiome composition.

### Thallus morphological type, reproduction, and secondary metabolites affect the photobiome and mycobiome

Lichen thallus morphology results from repeated, independent evolutionary innovations (Spribille et al. 2022). Foliose and fruticose thallus morphologies are thought to adapt to good hydration due to rain, dew, or vapor (Gauslaa 2014). In our dataset, crustose lichens consistently had a richer and more diverse photobiome, while foliose and fruticose lichens, even from the same mycobiont order, had a less diverse photobiome, with only one or two ASVs. Helms et al. (2001) consistently reported that crustose lichens in the Physciaceae are associated with a broader range of algae worldwide compared to other morphological types. Foliose and fruticose thalli have a complex, corticated structure and smaller contact with the substrate, which may limit access to environmental algae, while crustose thalli can encapsulate the substrate’s algae during their growth. However, it is unclear, whether this trend also results from more active physiological traits that may be selected for specific functional traits of the alga.

The lichen mycobiome has been described for lichens that are crustose (e.g., Muggia et al. 2016; Cometto et al. 2022), foliose (e.g., Muggia & Grube 2010; Tripathi et al. 2014) and fruticose (e.g., U’Ren et al. 2012; Spribille et al. 2016). However, to our best knowledge, these three morphologies have not been compared within similar (here, within Caliciales or Lecanorales) and different (between orders) evolutionary frames. In our study, the richness of the mycobiome of all three thallus morphological types was similar within one order while decreasing significantly in Pertusariales in comparison to other orders. However, within Lecanorales, the fruticose and crustose species had a more diverse mycobiome than foliose species, suggesting in this case that morphology allowing larger contact with the substrate (here, the bark of a tree) implies easier fungal colonization (as above for algae). Alternatively, since most fruticose Lecanorales considered in this study produce usnic acid derivatives and beta orcinol depsidones, in contrast to foliose and crustose Lecanorales, these secondary metabolites may change mycobiome diversity and composition. We were able to sample fruticose lichens from only two genera, *Usnea* and *Ramalina* (61 and 4 specimens, respectively), so a larger sampling may challenge our conclusions for Lecanorales. Furthermore, crustose Pertusariales had fewer OTUs and lower diversity than crustose species of other orders. This may result from their specific secondary metabolites (e.g., most of them contain xanthones and beta-orcinol depsidones). Each order of lichenized fungi tends to stabilize its mycobiome by means of particular secondary metabolites (see below), while the thallus morphological type does not play a major role in the acquisition of the mycobiome.

The influence of lichen reproductive strategies on the photobiome confirmed previous observations that lichens reproducing by spores and, therefore, recruiting new photobiome at settlement, associate with a broad range of algae (Wornik & Grube 2010; Cao et al. 2015; Steinová et al. 2019). Accordingly, lichen species that propagate via vegetative propagules display a narrower range of algae (Wornik & Grube 2010; Otálora et al. 2010). Conversely, the mycobiome did not display differences between sexually and asexually reproducing lichens for OTU richness, diversity, and community composition. Therefore, the mycobiome, in comparison to the photobiome of asexually reproducing lichens, probably undergoes different acquisition mechanisms similar to those of sexually reproducing lichens – mostly from the surrounding environment, although this requires further studies. Nevertheless, lichens with isidia have higher fungal OTU abundance and diversity than those with soredia. On the one hand, isidia have higher volumes, creating additional space for the transmission of fungi, while soredia that have no cortex are more open to fungal colonization during dispersion, but this seems not to compensate in terms of mycobiome diversity. Moreover, it has not been demonstrated that vertical transmission enriches the mycobiome of lichens with isidia.

Finally, lichen secondary metabolites structured the photobiome and mycobiome, suggesting one potential mechanism by which mycobiont identity influences the holobiont. On the one hand, the fact that the photobiome displays higher alpha diversity in lichens without any secondary metabolites suggests that these metabolites are somewhat restrictive on some algae (as stated by Băckor et al. 1998, 2010; Lokajová et al. 2014; Kosecka et al. 2022). However, we detected greater algal diversity in lichens with orcinol depsides, which are toxic to some algae (Bačkor et al. 1998, 2010; Lokajová et al. 2014). Either these depsides do not directly contact the algae within these lichens (here, we did not consider the localization of secondary metabolites, or their concentration in lichen thalli), or algae developed protective mechanisms against their phytotoxicity, at least at the symbiotic stage, during co-evolution (Lokajová et al. 2014). The presence of beta-orcinol depsides like atranorin and terpenoids like zeorin correlated with an increase of extraneous and endolichenic fungi, and thus the whole mycobiome, suggesting a limited effect on the mycobiome composition. The presence of orcinol depsides like evernic and lecanoric acids correlated with the increased number of lichenicolous fungi, suggesting tolerance or resistance of some of these parasites to such secondary metabolites. This supports previous observations that many lichen-associated fungi adapt to lichen secondary metabolites (Hawksworth 1982; Lawrey et al. 1999; Torzilli et al. 1999) and have roles other than simply antimicrobial (e.g., Goga et al. 2020; see Introduction). Yet, our data are correlative and do not directly estimate the impact of secondary metabolites on photobiome and mycobiome structure.

### Interaction network analysis gives clues to holobiont assembly

Low connectance and anti-nestedness are network features of highly intimate interaction (Guimarães Jr et al. 2007; Fontaine et al. 2011), as expected in the tight photobiont-mycobiont symbiosis. Furthermore, symbioses with strongly intimate interaction often show high specialization and high modularity in interaction networks (Thompson 2005; Pires & Guimarães Jr 2013; Guimarães Jr et al. 2017; Hembry et al. 2018). We observed here a modular and anti-nested network structure with low connectance between holobiont components for all the network linking (i) mycobiont and photobiome, (ii) mycobiont and mycobiome, and (iii) photobiome and mycobiome. However, the networks, including the mycobiome, revealed limited evidence of phylogenetic signals and low specialization levels in interaction with either the mycobionts or the photobiome. This contrasts with the frequent assumption of co-speciation between (i) lichens and lichenicolous fungi (but this co-speciation has rarely been analyzed from a macroevolutionary perspective; Werth et al. 2013; Diederich et al. 2018) and (ii) mycobiome and photobiome (for which the potential to build long-term, persisting interaction was suggested, but not further demonstrated; Muggia & Grube 2018).

In our study, the mycobiont significantly influenced the particular components of the mycobiome (lichenicolous and endolichenic fungi), but not the extraneous fungi. This may be due to the fact that the latter category is determined less by the mycobiont than by other factors such as the environment (in line with the above conclusion that the surrounding environment is a source of colonizing fungi). The lack of strong mycobiont-mycobiome and mycobiome-photobiome interactions may be due to our approach, which merges all fungi without consideration of their trophic mode in the assessment of networks. That is the most significant limitation in mycobiome studies (Muggia & Grube 2018), and our study is subject to the limitations of FUNGuild in addressing the trophic mode, even after careful expert visual correction. However, this convenient approach left 43% of identified OTUs trophically unassigned. This problem is prolonged by the scarcity of studies on lichen-associated fungi in general, and particularly in the studied geographical region, many of which are unculturable (Muggia & Grube 2018).

The interplay between the photobiome and mycobiome, estimated here for the first time to the best of our knowledge, appears low but significant. The finding of phylogenetic signals in all networks (strongest for the mycobiont—photobiome network) suggests that the holobiont is shaping the joint evolution of its members.

### Specialization and anthropogenic disturbance

This study shows how the lichen community changes in terms of photobiome and mycobiome on a narrow spatial scale. Lichen holobiont assembly depends mainly on the lichenized fungi – the mycobiont. Nevertheless, our study suggests a bigger specialization of the genus-specific preferences of the networks linking the mycobionts to the photobiome and to the mycobiome. Berlinches de Gea et al. (2022) showed that ecological fragmentation resulting from anthropogenic disturbance in Mediterranean forests decreases the specialization of mycobionts towards their algal partner in foliose and fruticose lichen with vegetative diaspores. We observed a generally increased specialization in the mycobiont-algae interaction for lichens with foliose thallus morphological type and with isidia or soredia. Foliose and fruticose (not crustose) lichens decreased their specialization under the influence of anthropogenic disturbances in the case of mycobiont-algae and mycobiont-mycobiome interaction. Furthermore, the ability of crustose lichens to encapsulate microbes during their growth may explain why they react less, although this requires further studies. We considered only two levels of anthropogenic disturbance, but together with Berlinches de Gea et al. (2022) we suggest a significant disruption of the composition of the holobiont, which may weaken the stability of the symbiosis in disturbed environments.

### Conclusion and perspectives

We attempted to uncover the processes governing lichen holobiont assembly. We revealed that the mycobiont – the main player in lichen symbiosis due to features such as thallus morphological type, reproductive strategies, and secondary metabolites – controls photobiome and mycobiome structure. This may be linked to the acquisition process (e.g., greater openness to the environment of the crustose thallus or vertical transmission by isidia and soredia), but other mechanisms with a co-evolutionary background may act in screening the partners. Indeed, phylogenetic signals occurred in the interaction networks between the partners of the lichen holobiont, especially between mycobionts and algae. Further studies focusing on evolutionary processes leading to morphological, anatomical, and chemical diversification of lichen thallus, combined with the consideration of biotic and abiotic factors, are needed to better understand the trends observed in this study.

We omitted bacteria and viruses, which are understudied parts of the lichen holobiont (Aschenbrenner et al. 2014; Ponsero et al. 2021). Indeed, many bacteria have been shown to be host-specific (Cardinale et al. 2012a, 2012b) and to change with photobiome shift (Xu et al. 2022). The interplay of these other players during lichen assembly also deserves attention in the future.

## Acknowledgments

We would like to thank Maciej Pazdro for the graphical support and David Marsh for editing the English. We greatly acknowledge members of Herbario Nacional de Bolivia, Instituto de Ecología, Universidad Mayor de San Andrés La Paz, for their generous cooperation and all the protected areas staff (http://sernap.gob.bo) for their help during the fieldwork. Lichens were collected in Bolivia with the permission of Ministerio de Medio Ambiente y Agua and in cooperation with Herbario Nacional de Bolivia (LPB).

## Author contributions

Conceptualization – MKo, AB, MK, BG-K, M-AS; Methodology – MKo, AB, BPL; Software – AB, BPL; Data curation – MKo, AB, BPL; Investigation – MKo, AB, BPL, MK; Validation – MKo, AB, BPL, BG-K, MK, M-AS; Formal analysis – MKo, AB, BPL; Supervision – MKo, AB, BG-K, MK, M-AS; Funding acquisition – MKo, BG-K, MK, M-AS; Visualization – MKo, AB, BPL; Project administration – MKo, AB; Resources – MKo, AB, BPL, BG-K, MK, AF, PR-F, M-AS; Writing - original draft – MKo, AB, BPL; Writing - review & editing – MKo, AB, BPL, BG-K, MK, AF, PR-F, M-AS.

## Data availability

Short read data generated in this study were deposited in the NCBI Sequence Read Archive with BioProject accession no. PRJNA1117370, which includes fungal and algal ITS2 rDNA sequencing data under BioSample accessions SAMN41564377 and SAMN41564378.

