## Supplementary material for "Factors shaping the assembly of lichen holobionts in a tropical lichen community": Supplementary_data_05_2024.pdf

### The following Supporting Information is available for this article:

**Fig. S1:** Rarefaction curves indicate that the sampled communities in La Paz are far from reaching saturation, while saturation is closer in Cochabamba, where we observed higher diversity in the undisturbed community (CB2) than in the disturbed one (CB1).
**Fig. S2:** The main mycobiont identity at the order level shapes both communities of the photobiome and mycobiome associated with lichens.
**Fig. S3:** The main mycobiont taxonomic assignment at the order (a) and the genus (b) level shapes the mycobiome associated with lichens.
**Fig. S4:** Variance partitioning indicated that the mycobionts and not the photobiome have the main impact on the mycobiome associated with lichens.
**Fig. S5:** Morphological type and reproductive strategies shape the photobiome and mycobiome associated with lichens.
**Fig. S6:** Reproductive strategies affect alpha-diversity of the photobiome and mycobiome associated with lichens.
**Fig. S7:** Human activities slightly influence the alpha-diversity of the photobiome and mycobiome associated with lichens.
**Fig. S8:** The type of network presents significantly different structures.
**Fig. S9:** The H2' index indicates that the strength and the significance of interaction specialization vary as a function of the type of network.
**Fig. S10:** The net relatedness index (NRI; which quantifies, for each OTU, the phylogenetic structure of its partner set based on mean pairwise phylogenetic distances) indicates that some mycobionts tend to associate with phylogenetically clustered partners (either the photobiome or mycobiome).
**Fig. S11:** The net relatedness index (NRI; which quantifies, for each OTU, the phylogenetic structure of its partner set based on mean pairwise phylogenetic distances) indicates that levels of specializations of mycobiome tend to be higher for endolichenic fungi compared with extraneous or lichenicolous fungi.
**Fig. S12:** The H2' index indicates that the strength of mycobiont-photobiome specialization is higher in foliose lichens with (at least partially) vegetative reproduction.
**Fig. S13:** Visualization of the different types of networks.

**Table S1:** Sampling localities.

**Table S2:** List of specimens used for analysis with taxonomic assignments of the mycobiont and information about thallus morphological type, reproductive strategies, and secondary metabolites.
**Table S3:** List of specimens used for analysis with taxonomic assignations of the mycobiont and information about thallus morphological type, reproductive strategies, and secondary metabolites.
**Table S4:** PERMANOVA revealed the significant effect of the main mycobiont taxonomy (genera) on the Bray-Curtis beta diversity of photobiome and mycobiome.
**Table S5:** Variance partitioning indicated that mycobionts, not the photobiome, have the main impact on the mycobiome, even at finer taxonomic levels.
**Table S6:** Lichens with similar secondary metabolites host similar photobiome and mycobiome.
**Table S7:** Phylogenetic signal in lichens.

**Methods S1:** Sample preparation, Polymerase Chain Reaction specification, and library preparation.
**Methods S2:** OTU table preparation.

**Supplementary Figures:**

**Fig. S1: Rarefaction curves indicate that the sampled communities in La Paz are far from reaching saturation, while saturation is closer in Cochabamba, where we observed higher diversity in the undisturbed community (CB2) than in the disturbed one (CB1).**

For mycobionts, photobiome, or mycobiome, the rarefactions plots indicated the mean number of fungal OTUs (OTU richness) or the mean phylogenetic diversity of the fungal OTUs (Faith's index) found in each community as a function of the number of sampled individual lichens.

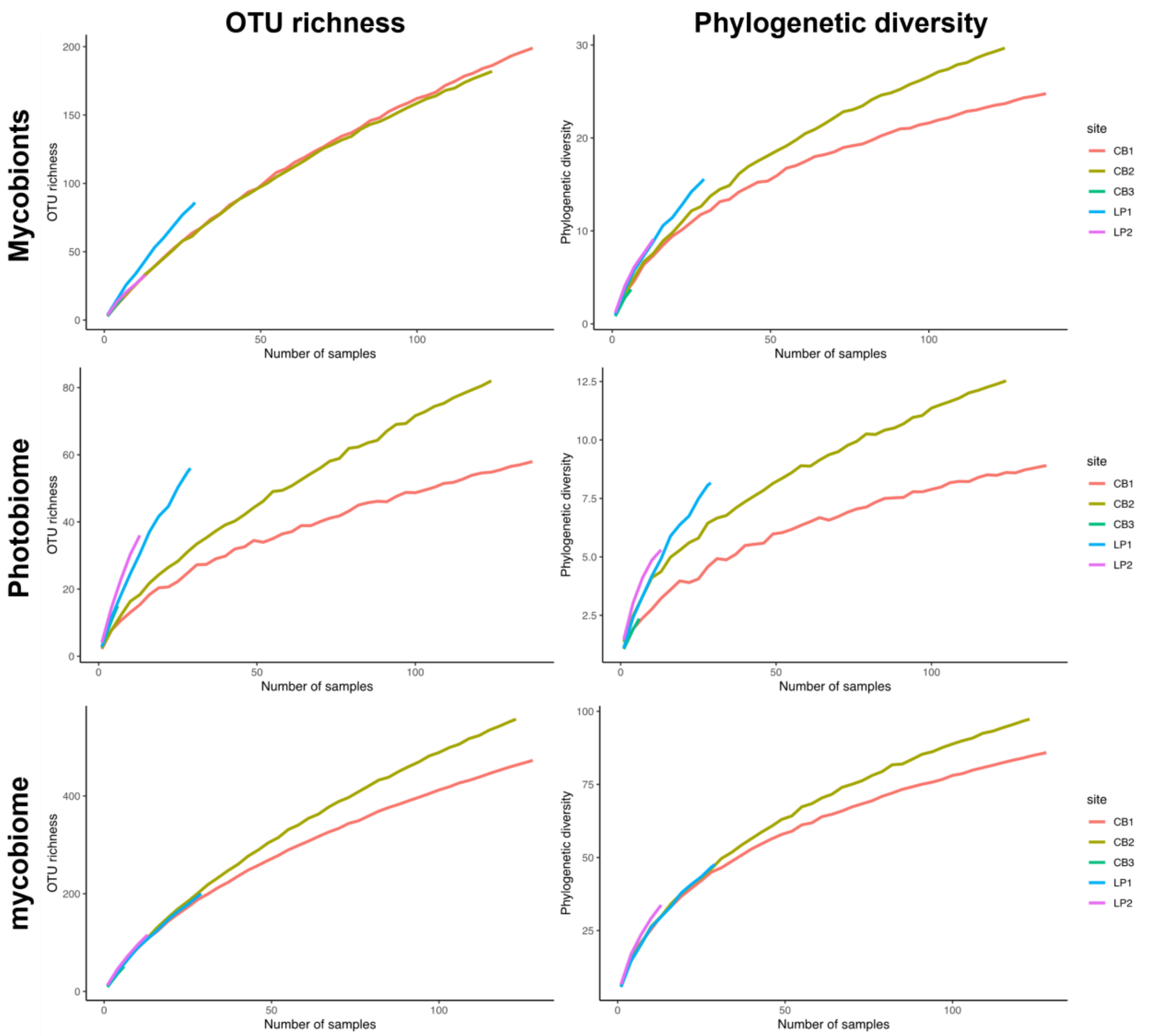


**Fig. S2: The main mycobiont identity at the order level shapes both communities of the photobiome and mycobiome associated with lichens.**

PCoA based on Bray-Curtis dissimilarity obtained from all specimens (n = 309). Shapes indicate the sample site (Cochabamba or La Paz), whereas colors present the main four mycobiont identity at the order level (a, c) (n = 309). PERMANOVA test revealed significant differences within mycobiont orders in both photobiome ($R^2$=0.11, p=0.001) and mycobiome ($R^2$=0.05, p=0.001). Relative abundances of the most abundant algal (b) and fungal (d) groups at the species and order levels, respectively.

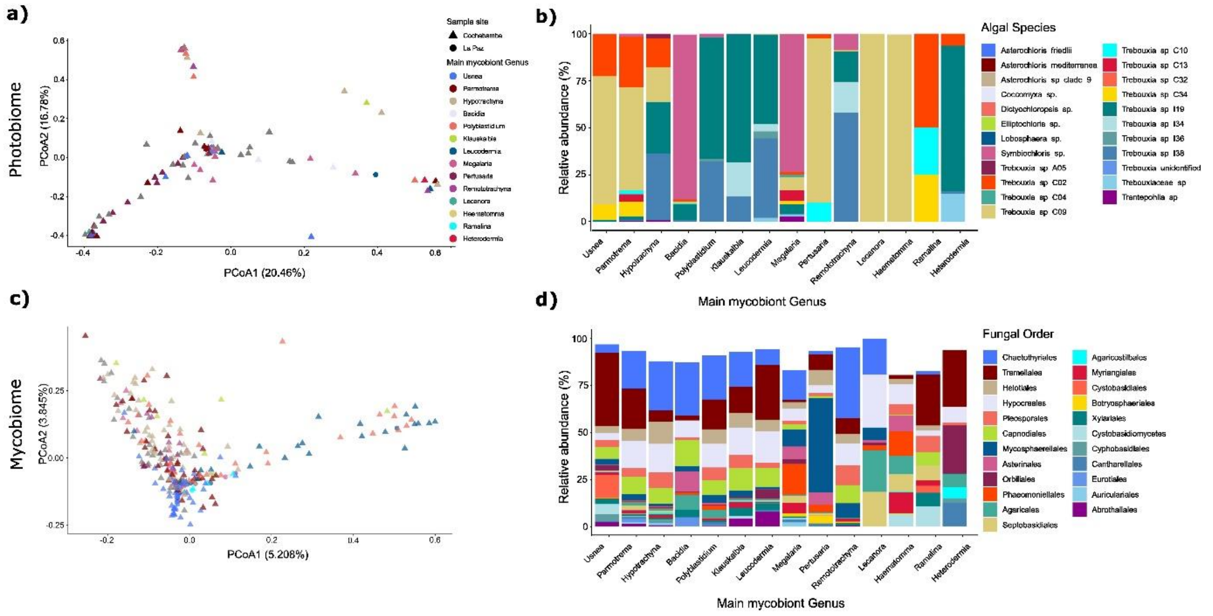


**Fig. S3: The main mycobiont taxonomic assignment at the order (a) and the genus (b) level shapes the mycobiome associated with lichens.**

Relative abundances (RAs) of the trophic mode defined by using FunGuild database, in the total fungal community. After the verification, e.g., removing animal or plant pathotrophs to extraneous fungi, pathotrophs, symbiotrophs, and saprotrophs correspond respectively to lichenicolous, endolichenic, and extraneous fungi.

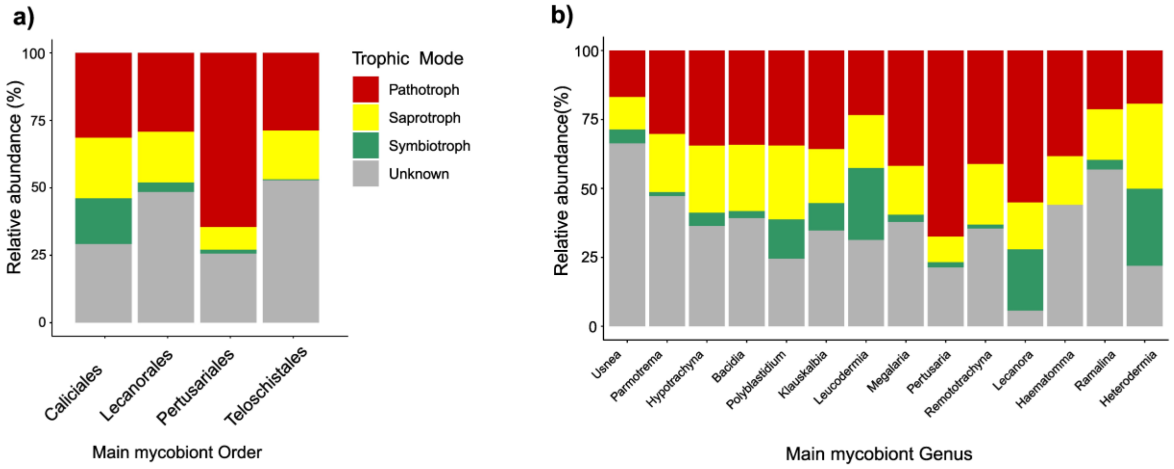


**Fig. S4: Variance partitioning indicated that the mycobionts and not the photobiome have the main impact on the mycobiome associated with lichens.**

For all sites or each site separately, we partitioned the variance between mycobiome (measured using Bray-Curtis dissimilarities or weighted UniFrac distances) as a function of the distances in mycobiont or photobiome communities (also measured using Bray-Curtis dissimilarities or weighted UniFrac distances). Using Venn diagrams, we reported the percentage of variance that is explained by the mycobionts or the algal communities or by a combined effect of both communities.

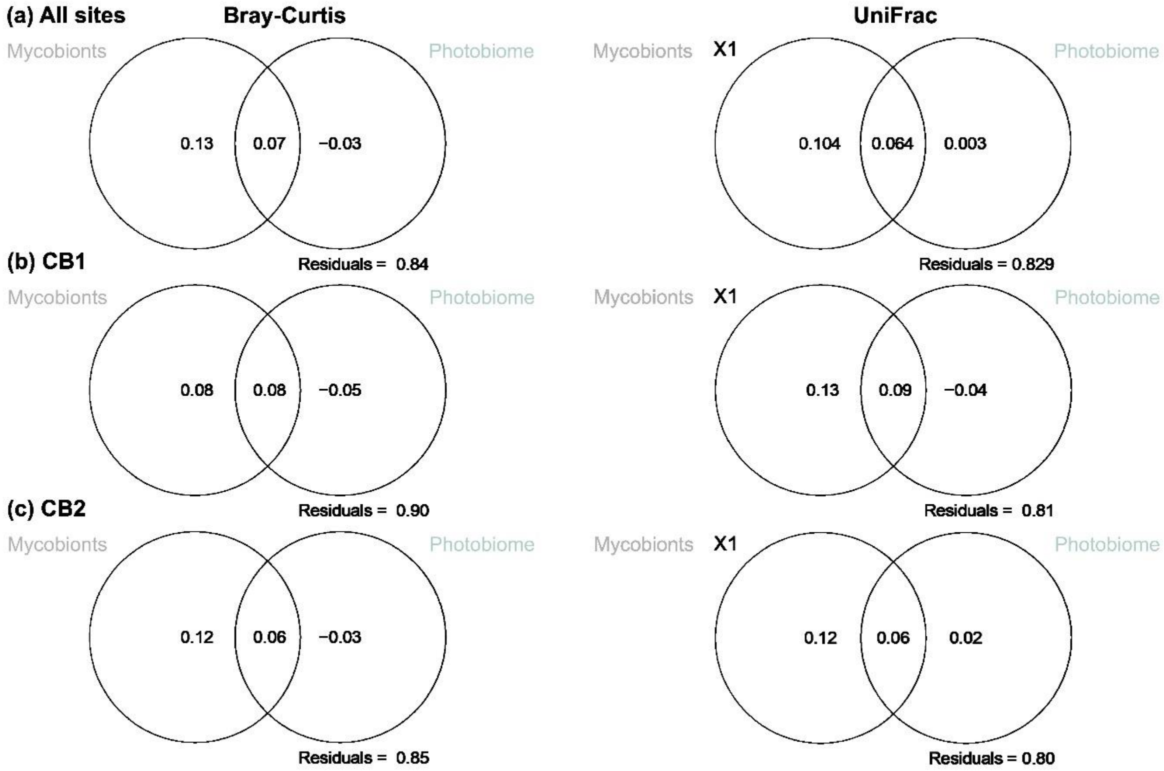
(a) All sites
Bray-Curtis
UniFrac
Mycobionts
Photobiome
0.13
0.07
−0.03
Residuals = 0.84
Mycobionts
X1
Photobiome
0.104
0.064
0.003
Residuals = 0.829
(b) CB1
Mycobionts
Photobiome
0.08
0.08
−0.05
Residuals = 0.90
Mycobionts
X1
Photobiome
0.13
0.09
−0.04
Residuals = 0.81
(c) CB2
Mycobionts
Photobiome
0.12
0.06
−0.03
Residuals = 0.85
Mycobionts
X1
Photobiome
0.12
0.06
0.02
Residuals = 0.80

**Fig. S5: Morphological type and reproductive strategies shape the photobiome and mycobiome associated with lichens.**

Alpha-diversity (Shannon index) of samples depends on the combined effect of morphological type and main mycobiont taxonomic assignation (A, B); and on lichen reproductive strategies (C, D). A Kruskal-Wallis test ($P < 0.05$) was used for statistical analysis. Letters indicate significant differences between conditions.

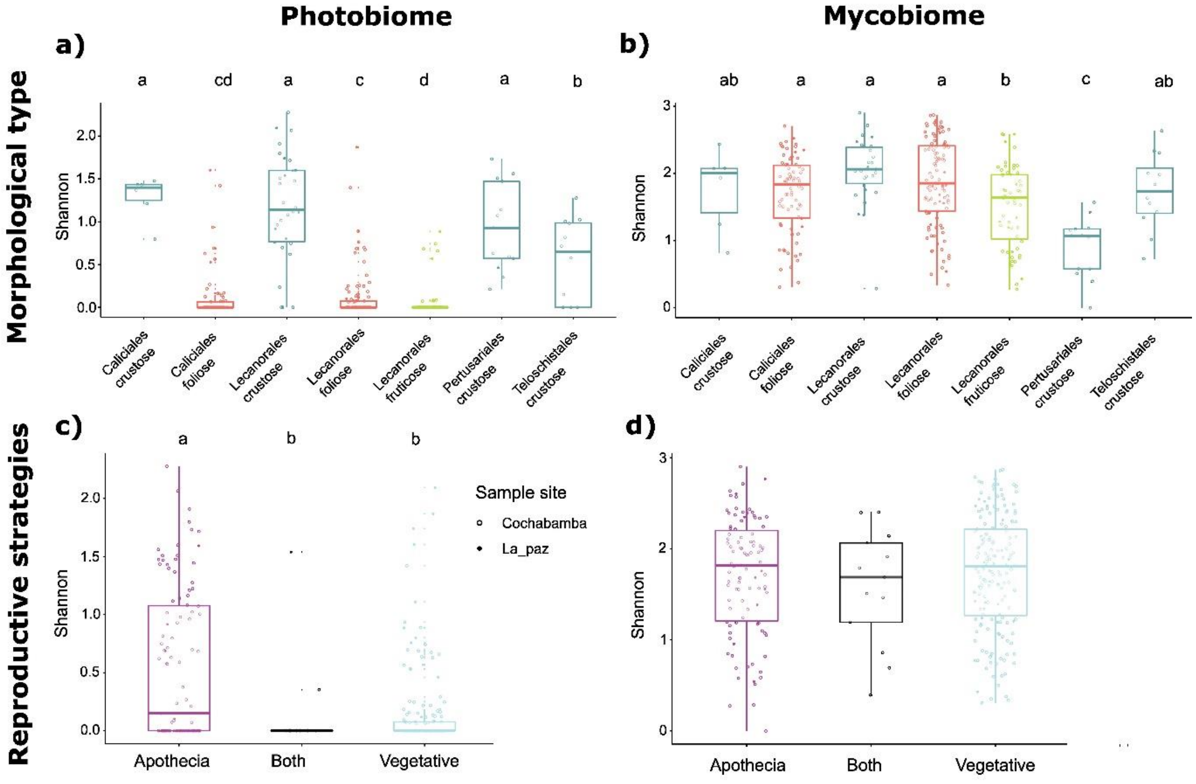

**Fig. S6: Reproductive strategies affect alpha-diversity of the photobiome and mycobiome associated with lichens.**

Alpha-diversity (observed fungal OTUs and algal ASVs, and Shannon index) of samples depending on reproductive strategy (A: apothecia, AVI: apothecia and isidia, AVIVS: apothecia, isidia and soredia, AVS: apothecia and soredia, ST: sterile, VI: isidia, VIVS: isidia and soredia, VS: soredia). A Kruskal-Wallis test ($P < 0.05$) was used for statistical analysis. Letters indicate significant differences between conditions.

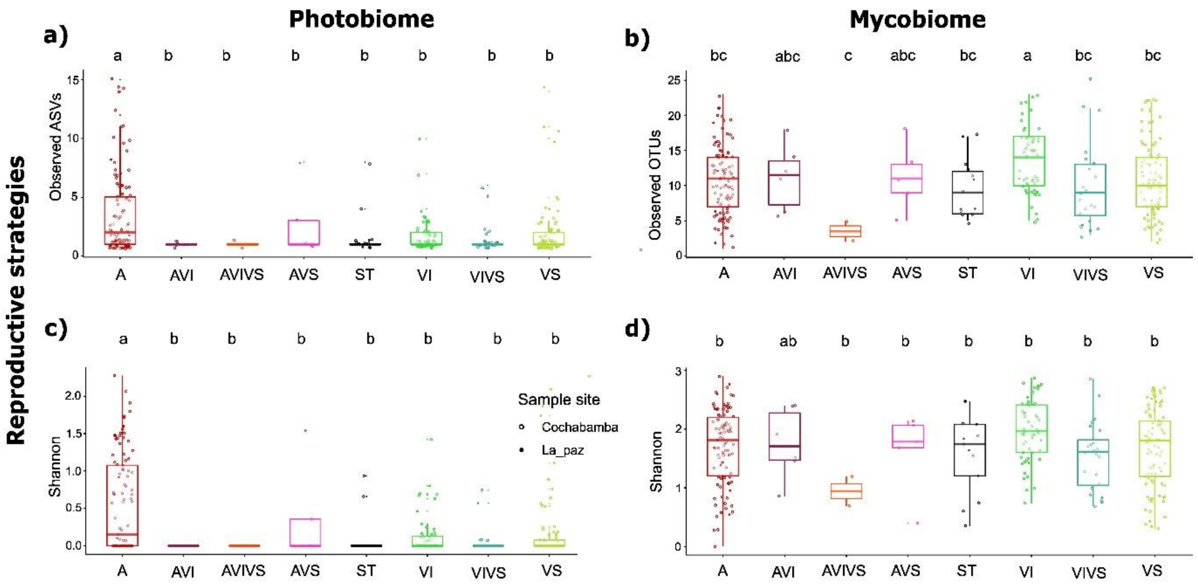

**Fig. S7: Human activities slightly influence the alpha-diversity of the photobiome and mycobiome associated with lichens.**

Relative abundances (RAs) of the most abundant mycobionts (a), algae (b), and mycobiome (c) at the genus, species, and order levels, respectively. Alpha-diversity (observed fungal OTUs and algal ASVs (d, e, f), and Shannon index (g, h, i) of samples depending on localities (LP1, LP2, CB1, CB2, and CB3). A Kruskal-Wallis test ($P < 0.05$) was used for statistical analysis. Letters indicate significant differences between conditions. Mycobiome corresponds to the total fungal community except for the main mycobiont and additional mycobionts.

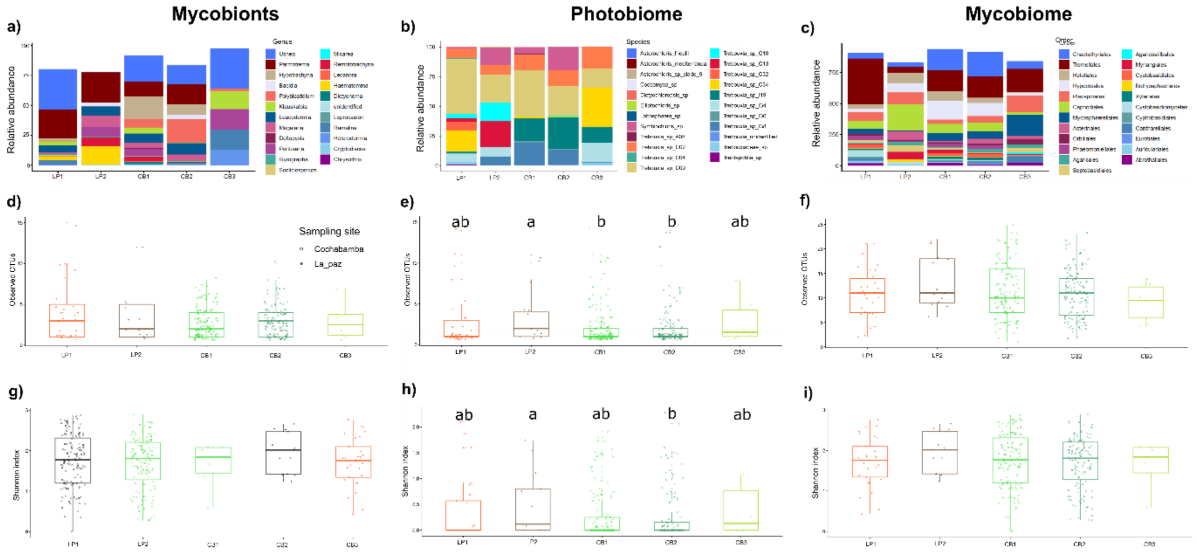

**Fig. S8: The type of network presents significantly different structures.**

For each site and each type of network, we measured the original connectance, modularity, and nestedness values (represented in red) that we compared to the null expectations given by randomized networks (in gray). Dashed lines delimit the estimated 95% confidence intervals.

a) Networks between mycobionts and photobiome

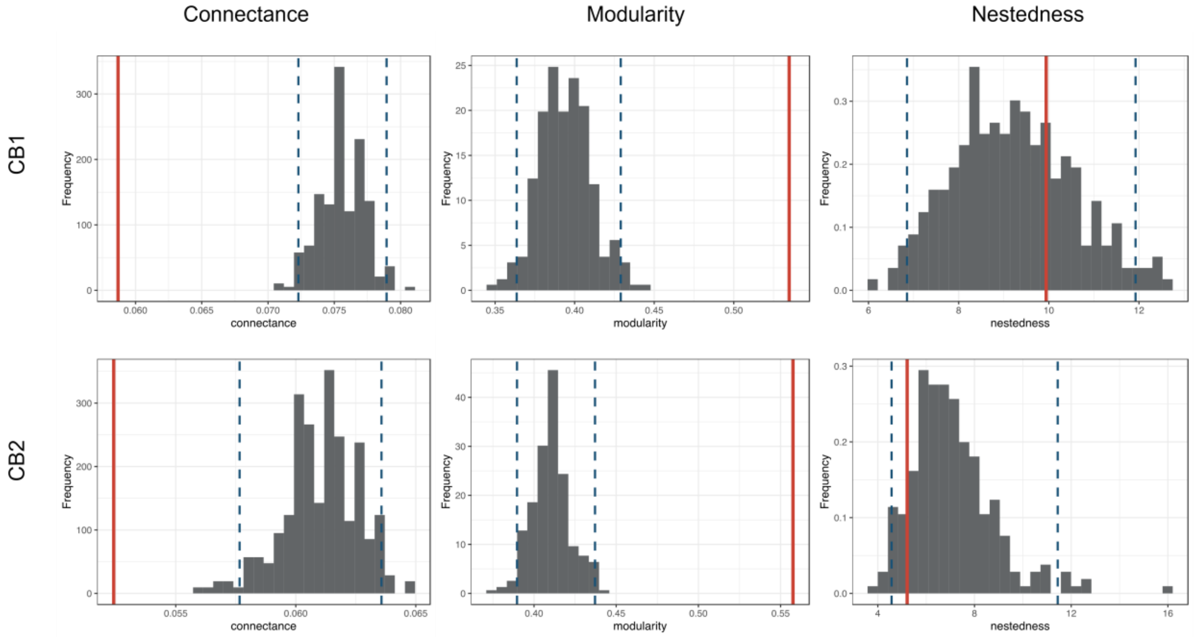


b) Networks between mycobionts and mycobiome

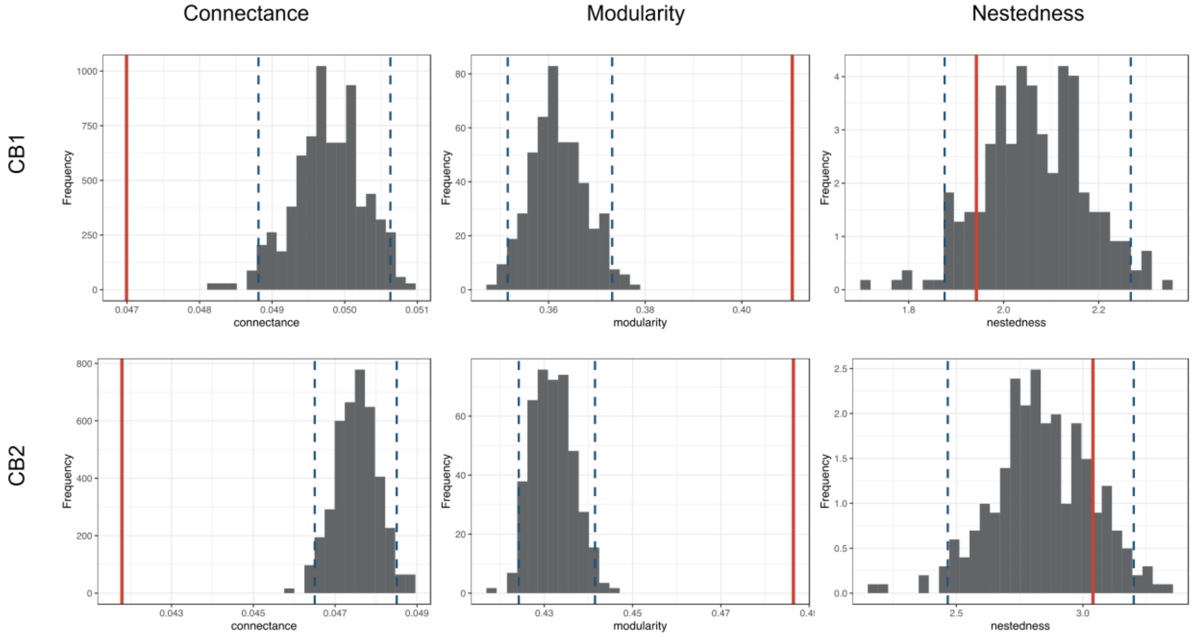

c) Networks between photobiome and mycobiome

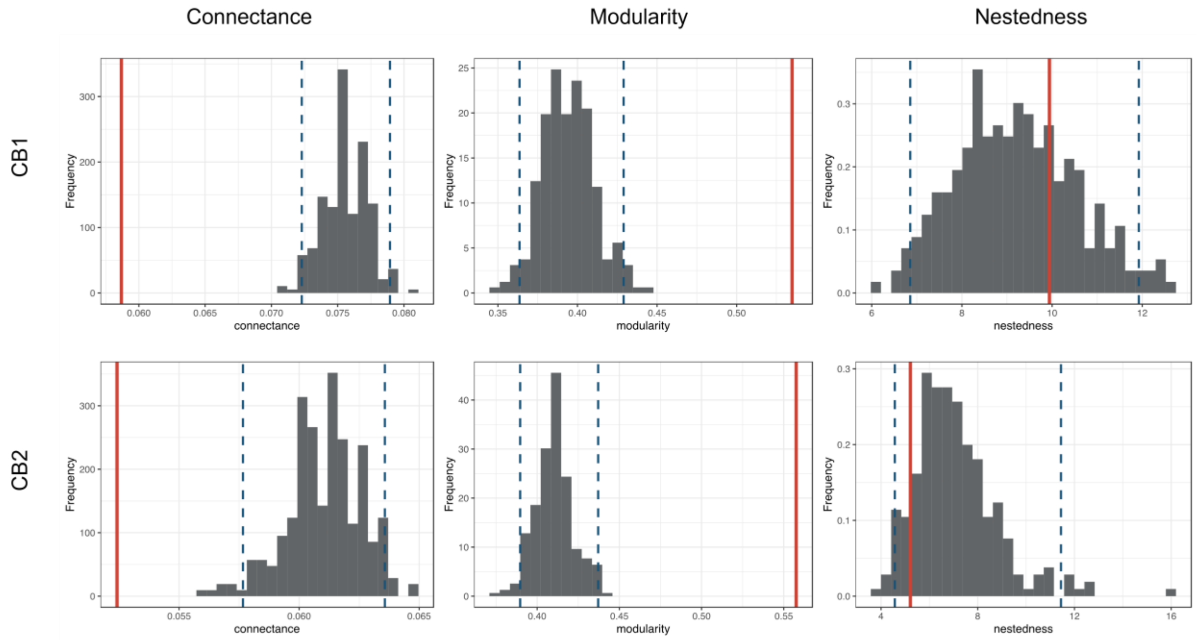


**Fig. S9: The H2' index indicates that the strength and the significance of interaction specialization vary as a function of the type of network.**

For each site and each type of network, we measured the original H2' value (represented in red) that we compared to null expectations given by randomized networks (in gray). Dashed lines delimit the estimated 95% confidence intervals. An H2' value close to 0 indicates low specialization, whereas H2' value close to 1 indicates high specialization.

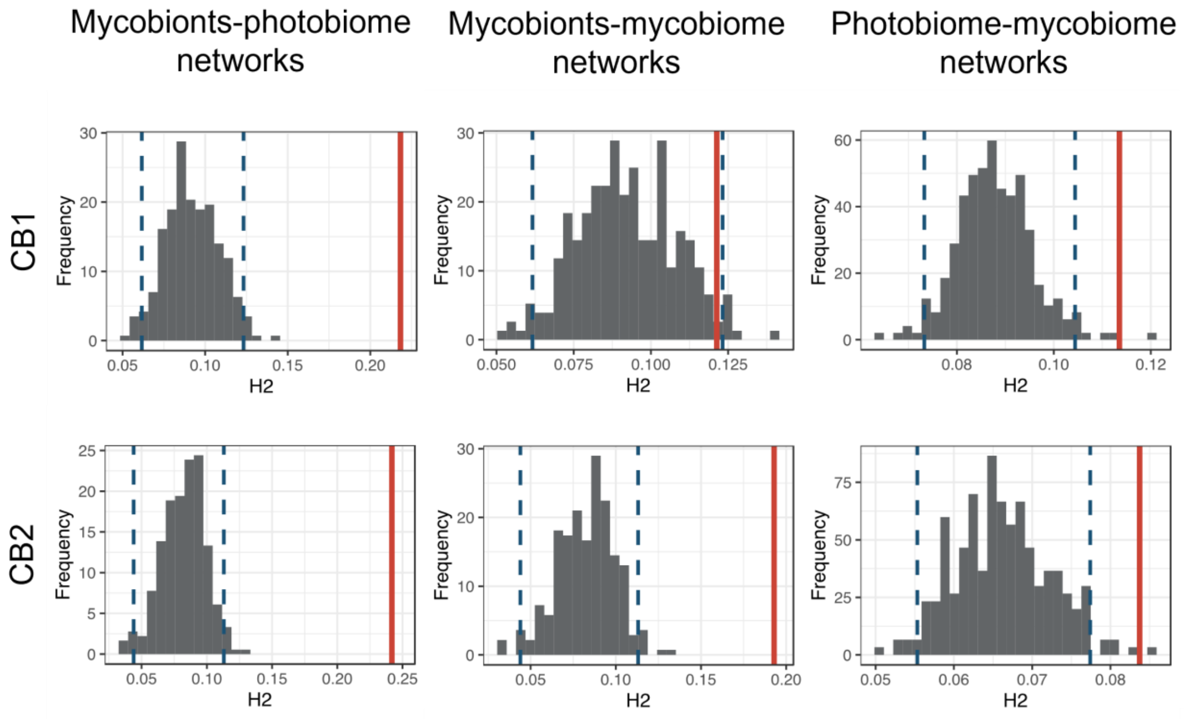

**Fig. S10: The net relatedness index (NRI; which quantifies, for each OTU, the phylogenetic structure of its partner set based on mean pairwise phylogenetic distances) indicates that some mycobionts tend to associate with phylogenetically clustered partners (either the photobiome or mycobiome).**

For each site, we measured - in the (a) mycobionts-photobiome network, (b) the mycobionts-mycobiome network, or (c) the photobiome-mycobiome network - the NRI values for each OTU and assessed its significance using permutations. We reported here the NRI values as a function of the mycobiont genera, algal species, or mycobiome-forming fungi orders. Significant positive NRI values are indicated in green (i.e. phylogenetic clustering of the partners), whereas non-significant NRI values are in purple. We tested the effect of the taxonomic groups on the NRI values using a Kruskal-Wallis test (reported above each panel). Results were qualitatively similar when using PSS (instead of NRI).

**(a) mycobionts-photobiome network**

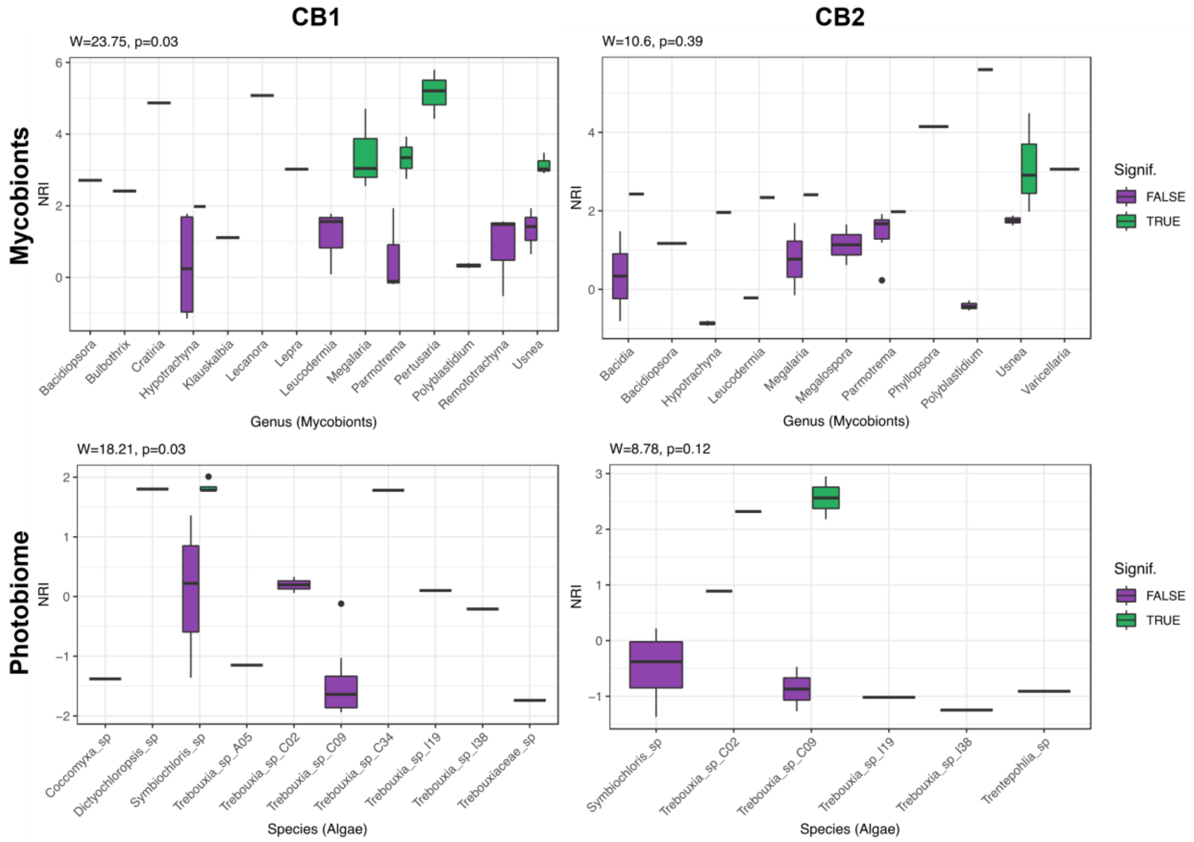

### (b) mycobionts-mycobiome network

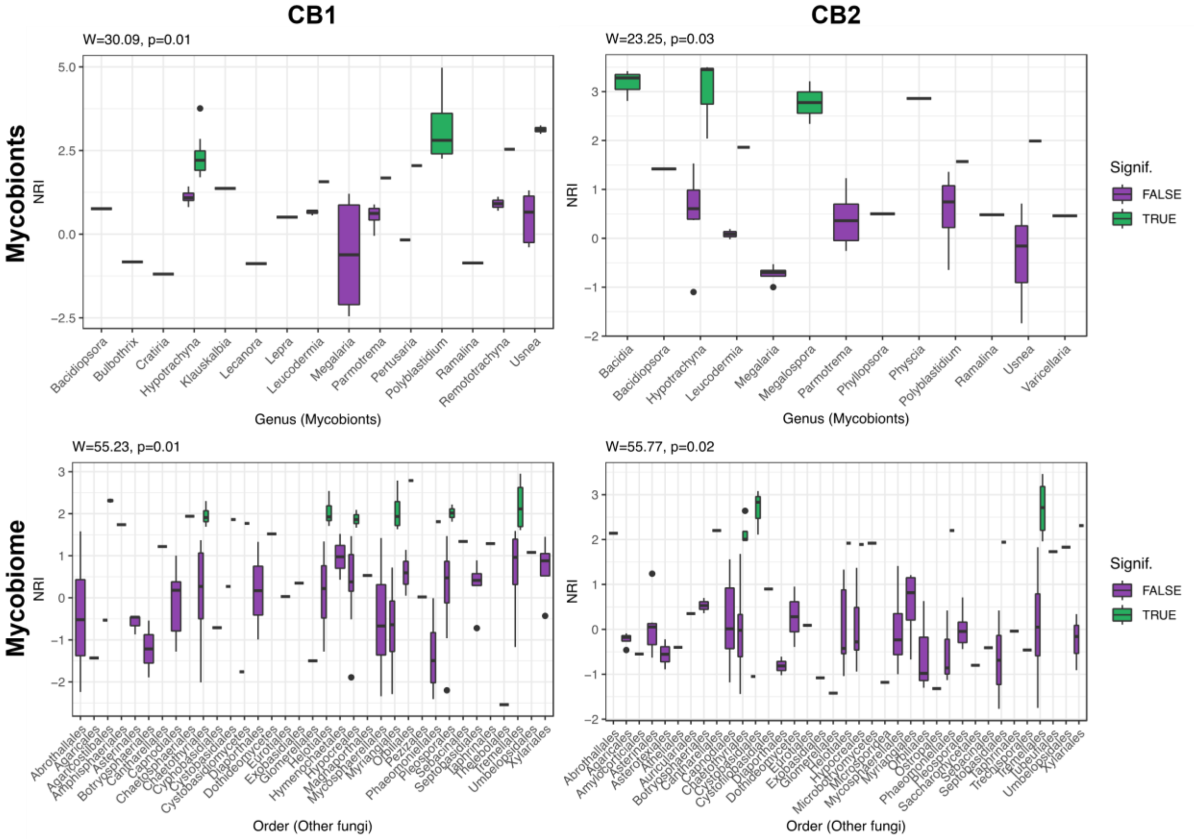

### (c) photobiome-mycobiome network

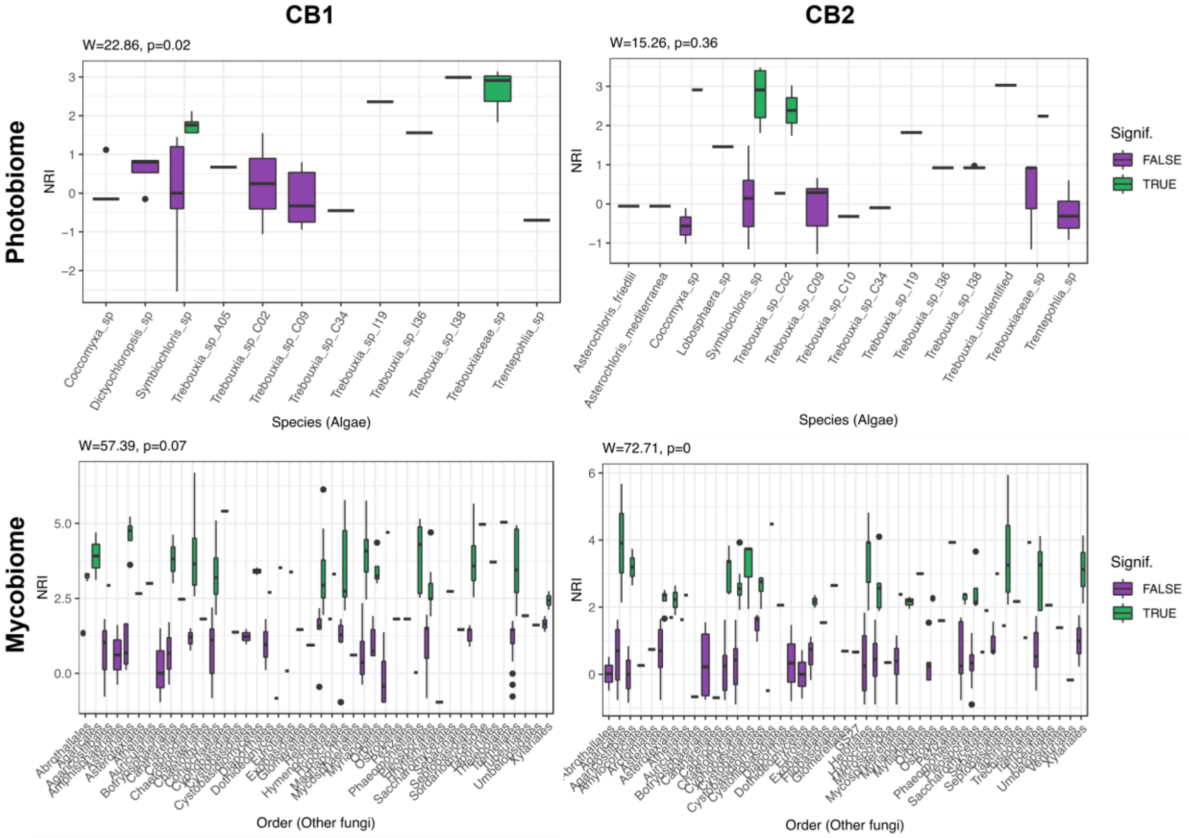

**Fig. S11: The net relatedness index (NRI; which quantifies, for each OTU, the phylogenetic structure of its partner set based on mean pairwise phylogenetic distances) indicates that levels of specializations of mycobiome tend to be higher for endolichenic fungi compared with extraneous or lichenicolous fungi:**

For each site, we measured - in (a) the mycobionts-mycobiome network or (b) the photobiome-mycobiome network - the NRI values for each OTU and assessed its significance using permutations. We reported here the NRI values as a function of the ecological guild inferred using FUNGuild. Significant positive NRI values are indicated in green (i.e. phylogenetic clustering of the partners), whereas non-significant NRI values are in purple. We tested the effect of the guilds on the NRI values using a Kruskal-Wallis test (reported above each panel).
Results were qualitatively similar when using PSS (instead of NRI).

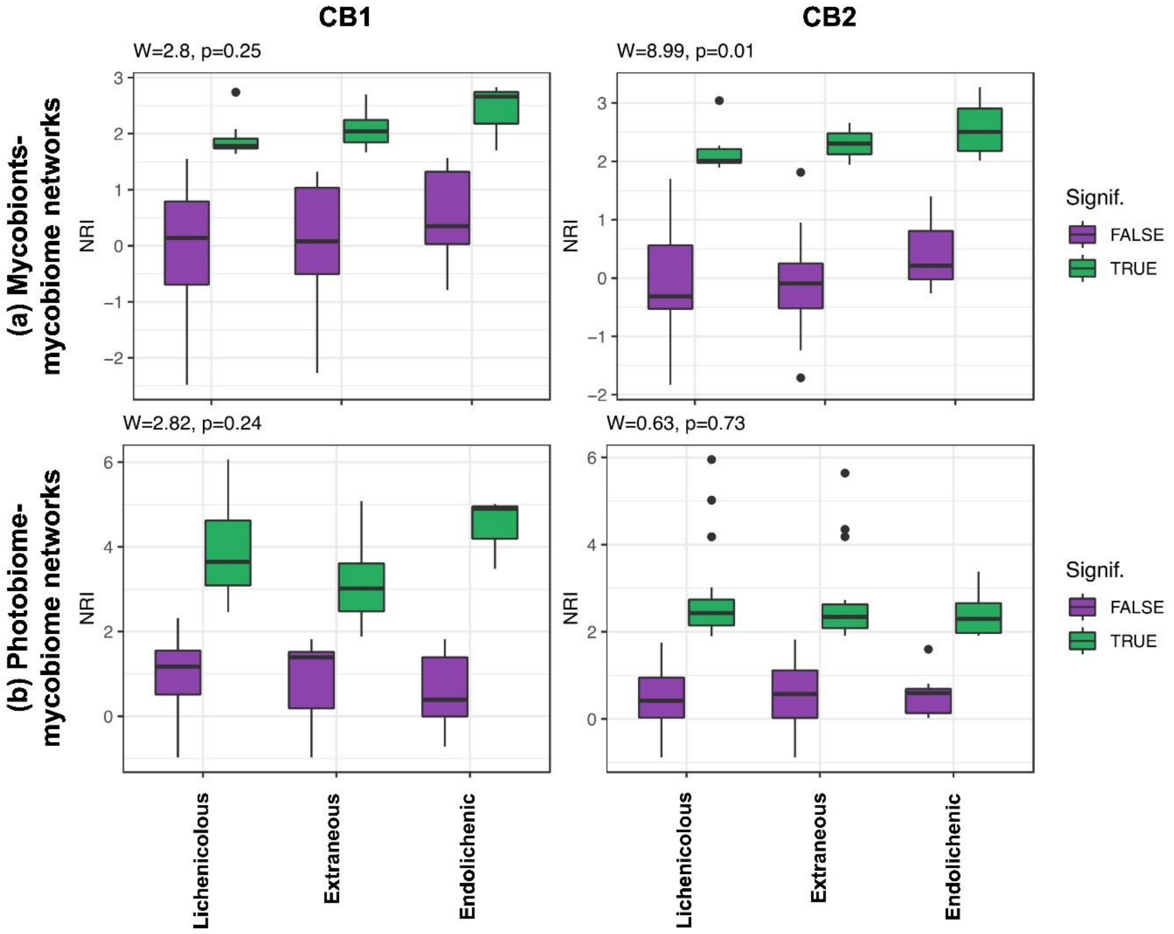

**Fig. S12: The H2' index indicates that the strength of mycobiont-photobiome specialization is higher in foliose lichens with (at least partially) vegetative reproduction.**

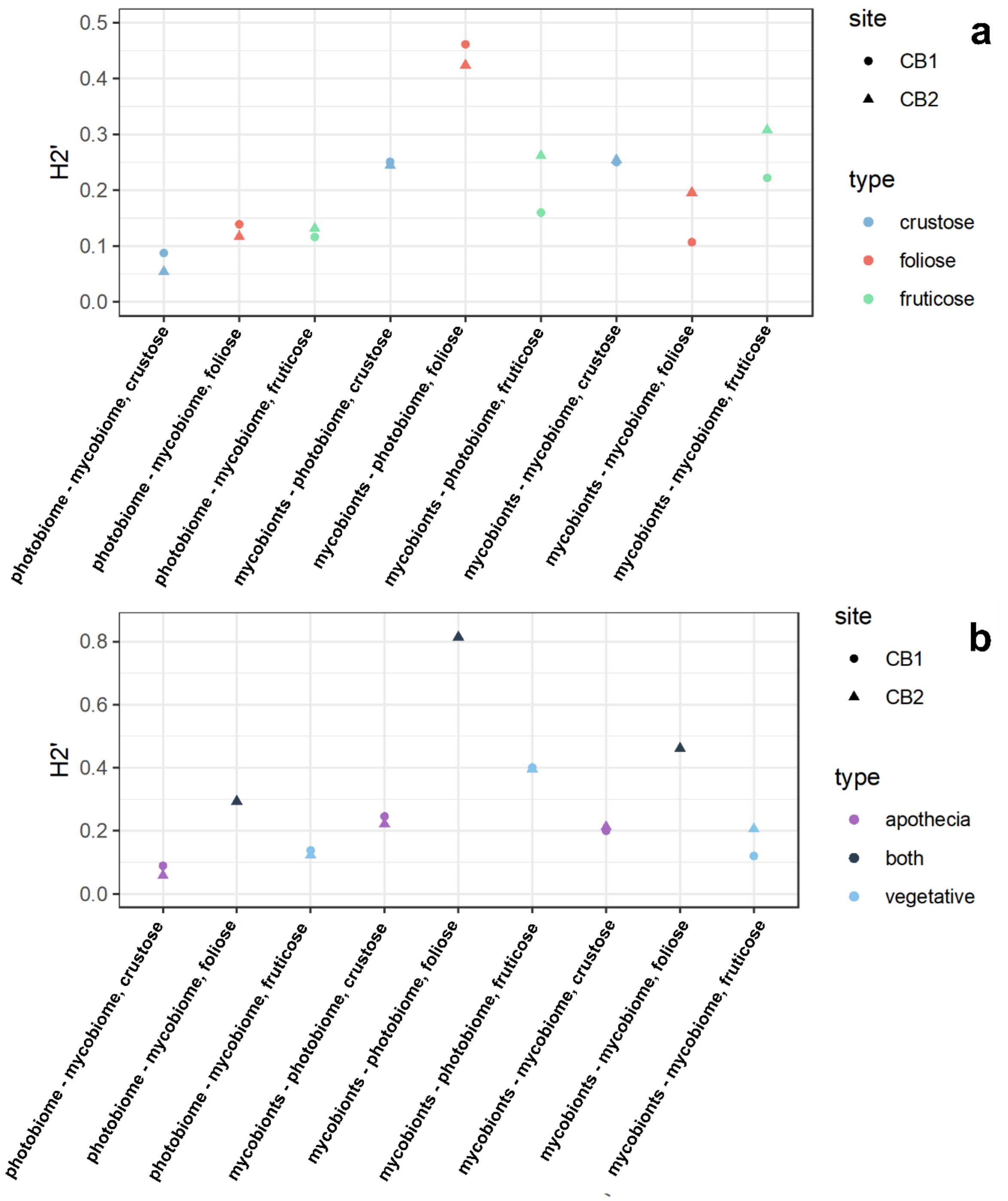

**Fig. S13: Visualization of the different types of networks.**

Each node represents an OTU (either a mycobiont, the photobiome, or the mycobiome) and each gray line indicates that there exists an interaction between two OTUs and their widths are proportional to the number of times the interaction has been observed. The position of the nodes reflects the similarity in species interactions using the Fruchterman-Reingold layout algorithm from the *igraph* R-package.

In the case of comparison between disturbed (CB1) and undisturbed (CB2) communities, the shapes are slightly different, especially for the network of mycobionts and the mycobiome (b), which reflects a decrease in specialization level.

a) Networks between mycobionts and photobiome

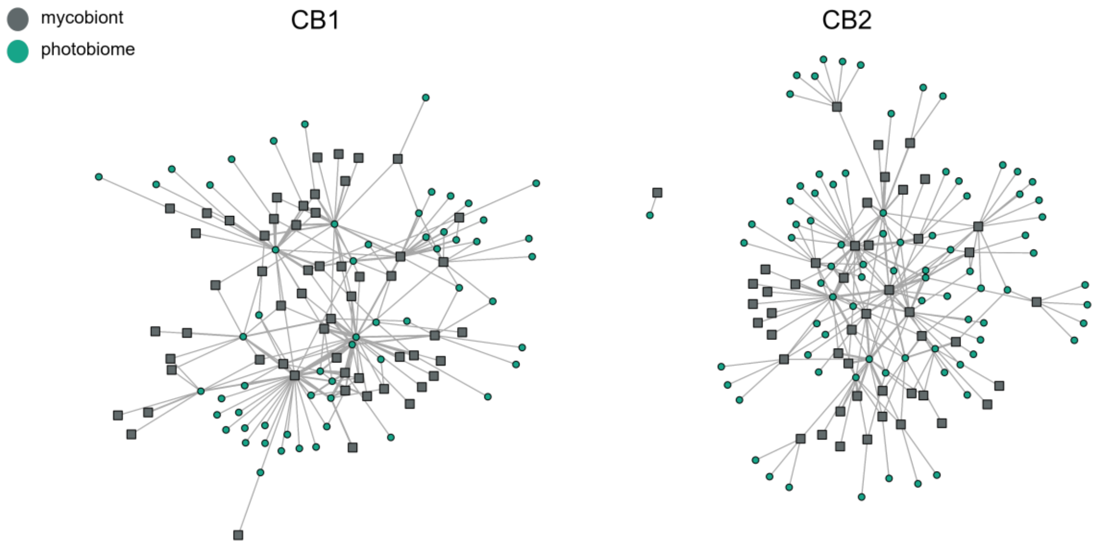

b) Networks between mycobionts and mycobiome

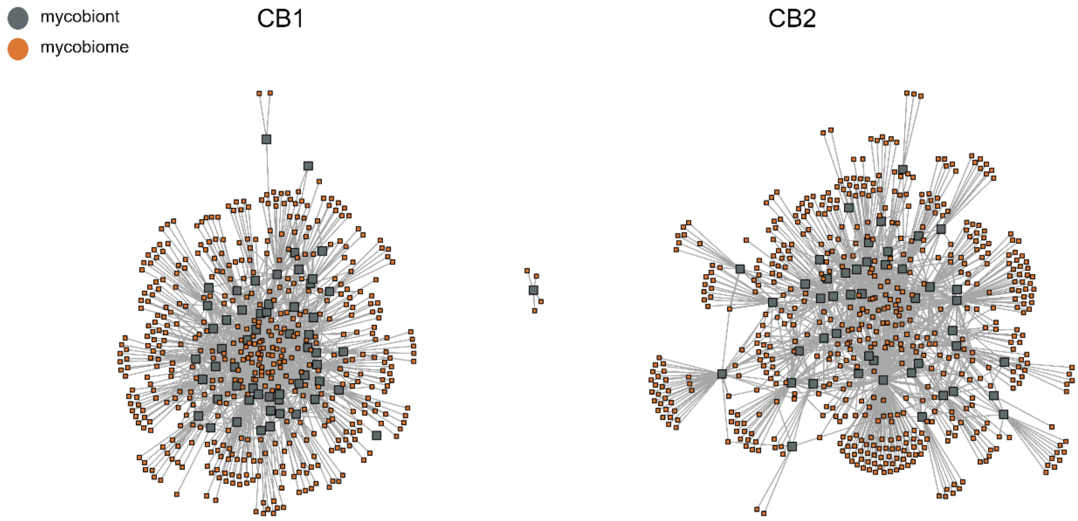


c) Networks between photobiome and mycobiome

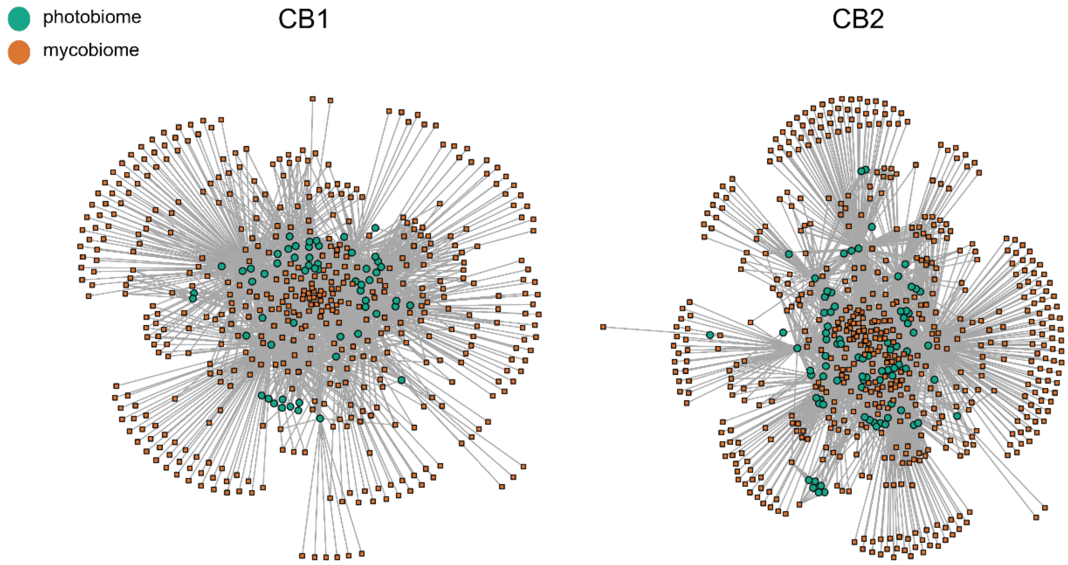

### Supplementary Tables

#### Table S1: Sampling localities.

Materials have been collected and deposited in the lichen collections of the University of Gdańsk (UGDA), Institute of Botany of Polish Academy of Sciences (KRAM) and Herbario Nacional de Bolivia, Universidad Mayor de San Andrés and Museo Nacional de Historia Natural La Paz (LPB) in 2016. The specimens have been collected by all rules regarding protected areas after receiving the relevant permissions. Material collection has been performed in cooperation with Universidad Mayor de San Andrés, La Paz.

The similar altitude, the same type of forest (Yungas cloud forest), and the proximity of localities may allow the influence of elevation above sea level and different climatic conditions on the mycobiome and photobiont pools to be excluded.

#### Table S2: List of specimens used for analysis with taxonomic assignments of the mycobiont and information about thallus morphological type, reproductive strategies, and secondary metabolites.

#### Table S3: List of orders and genera of lichen-forming fungi (mycobionts) identified in the fungal database accepted as main mycobionts according to the highest abundance and additional mycobionts.

#### Table S4: PERMANOVA revealed the significant effect of the main mycobiont identity (genera) on the Bray-Curtis beta diversity of photobiome and mycobiome.

To assess the effect of available indicators on the photobiome and mycobiome, we performed a permutational analysis of variance (PERMANOVA) (Excel sheet 1). We tested the effect of the sample site, habitat, reproductive strategies, morphological type, order, and genus of the main mycobiont. The results of the PERMANOVA (R2 and p-value based on 1,000 permutations) are indicated for all three communities. Significant correlations are highlighted in bold.

All factors are statistically significant ($p < 0.05$). However, most do not show the correlation (R2 < 0.05). We observed the most important effect of the genus of the main mycobiont for all communities. In the case of the photobiome, we also observe little effect on reproduction strategy, morphological type, and order of main mycobiont.

#### Table S5: Variance partitioning indicated that mycobionts, not the photobiome, have the main impact on the mycobiome, even at finer taxonomic levels.

For all sites or each site separately, we partitioned the variance between the mycobiome (measured using Bray-Curtis dissimilarities) as a function of the distances in the mycobiont or photobiome communities (also measured using Bray-Curtis dissimilarities). We performed the partitioning when looking at all lichens or when looking separately at the most abundant mycobiont genera. Here, we reported the effects of the photobiome and the mycobionts, which were assessed using an ANOVA (with 999 permutations). Significant contributions are highlighted in bold. Results were qualitatively similar when using weighted UniFrac distances.

**Table S6: Lichens with similar secondary metabolites host similar photobiome and mycobiome.**

For each site, we measured for each guild, using a Mantel test, whether lichens with similar secondary metabolites tend to host similar fungal (mycobiome or other mycobionts) and algal communities. For each test, we reported the Spearman correlation (between the “chemical distances” and the weighted UniFrac distances) and the corresponding p-value evaluated using 10,000 permutations.

**Table S7: Phylogenetic signal in lichen’s networks.**

For each type of network and each site, we measured each in guild phylogenetic signal in species interactions using a Mantel test. For each test, we reported the Spearman correlation (between the phylogenetic distances and beta diversities - measured using the weighted Jaccard or UniFrac distances) and the corresponding p-value evaluated using 10,000 permutations.

### Supplementary Methods:

#### Methods S1: Sample preparation, polymerase chain reaction specification, and library preparation.

Total lichen DNA was extracted using the Plant & Fungi DNA Purification Kit (Eurx, Poland). Before the triplicate by polymerase chain reaction (PCR), the concentration of each DNA isolate was measured by fluorescence (Quant-IT PicoGreen; Invitrogen, USA) and then diluted to a final 3.5 ng/μL. Fungal ITS2 rDNA was amplified using ITS86-F/ITS4 (White et al., 1990; Turenne et al., 1999) primer pair, while algal ITS2 rDNA by FDGITS2-f/FDGITS2-r (Dal Grande et al., 2018). Multiplexing was made by the use of a unique pair of barcoded primers in the PCR following Petrolli et al. (2021).

Polymerase chain reaction was performed with 25 μl of reaction mixture containing 0.2 mM of dNTPs, 0.3 mM of each primer, 2 units of DFS-Taq DNA Polymerase (BIORON, Germany), 1× Incomplete Buffer supplied with $MgCl_2$, BSA 3% and 3 μl template DNA. In the case of fungal communities amplification, the following program was used: initial denaturation, 10 min at 95°C; 30 cycles of denaturation, 30 s at 95°C; annealing, 30 s at 56.5°C; and elongation, 30 s at 72°C – final elongation 7 min at 72°C. Amplification of algal communities was performed with the following cycle conditions: initial denaturation, 4 min at 95°C; 35 cycles of denaturation, 30 s at 95°C; annealing, 20 s at 54°C; and elongation 20 s at 72°C – final elongation 5 min at 72°C.

Triplicate PCR reactions were pooled, and the quality of amplification was checked on agarose gels. Next, the amplicons were purified by Ampure XP beads (Agencourt, Beckman Coulter, USA). The concentration of PCR products was measured by fluorescence, then pooled together in equimolar amounts to build one library per group (fungi and algae). Each library was purified twice by Ampure XP beads and quantified by fluorescence. Sequencing was proceeded using a MiSeq, paired-end 2*250bp, Illumina technology (FASTERIS, Switzerland). Two negative controls (ultrapure water) were used per PCR trial (plate), resulting in a total of 8 negative controls per library.

**Methods S2: OTU table preparation.**

For the fungal dataset, we extracted the OTUs corresponding to lichenized fungi from each sample. We separated the most abundant lichenized fungi OTUs (referred to as the “main mycobiont”) from the additional lichenized fungi present in samples (Table S2). We checked that the taxonomic assignations of the main mycobiont identified using metabarcoding matched those based on morphology. We only kept samples with matching taxonomic assignations (Table S3).

The few samples having fewer than 1000 algal reads or less than 100 mycobiome reads were discarded in the following analyses. It resulted in a total of 309 samples (Table S3). We next reconstructed a phylogenetic tree of (i) all the lichenized fungi OTUs, (ii) all mycobiome OTUs, and (iii) all algal OTUs following Perez-Lamarque, Petrolli et al. (2022). For the algae dataset, we revised the taxonomy assignation by phylogenetic analysis and manually changed the assignation following Muggia et al. (2020), Medeiros et al. (2021), and Kosecka et al. (2022) in the case of the genus *Trebouxia* or keeping the highest confirmed classification of the others.
